# Functional roles of multiple Ton complex genes in a *Sphingobium* degrader of lignin-derived aromatic compounds

**DOI:** 10.1101/2021.08.06.455174

**Authors:** Masaya Fujita, Shodai Yano, Koki Shibata, Mizuki Kondo, Shojiro Hishiyama, Naofumi Kamimura, Eiji Masai

## Abstract

TonB-dependent transporters (TBDTs) mediate outer membrane transport of nutrients using the energy derived from proton motive force transmitted from the TonB−ExbB−ExbD complex localized in the inner membrane. Recently, we discovered *ddvT* encoding a TBDT responsible for the uptake of a 5,5-type lignin-derived dimer in *Sphingobium* sp. strain SYK-6. Furthermore, overexpression of *ddvT* in an SYK-6-derivative strain enhanced its uptake capacity, improving the rate of platform chemical production. Thus, understanding the uptake system of lignin-derived aromatics is fundamental for microbial conversion-based lignin valorization. Here we examined whether multiple *tonB*-, *exbB*-, and *exbD*-like genes in SYK-6 contribute to the outer membrane transport of lignin-derived aromatics. The disruption of *tonB2*−*6* and *exbB3* did not reduce the capacity of SYK-6 to convert or grow on lignin-derived aromatics. In contrast, the introduction of the *tonB1*−*exbB1*−*exbD1*−*exbD2* operon genes into SYK-6, which could not be disrupted, promoted the conversion of β-O-4-, β-5-, β-1-, β-β-, and 5,5-type dimers and monomers, such as ferulate, vanillate, syringate, and protocatechuate. These results suggest that TonB-dependent uptake involving the *tonB1* operon genes is responsible for the outer membrane transport of the above aromatics. Additionally, *exbB2*/*tolQ* and *exbD3*/*tolR* were suggested to constitute the Tol-Pal system that maintains the outer membrane integrity.

## Introduction

Lignin, a major component of plant cell walls, is the most abundant aromatic compound on Earth and is expected to be a renewable alternative to fossil resources^1,2^. Recently, the production of value-added chemicals from lignin by combining chemical depolymerization of lignin with bacterial catabolic systems for low-molecular-weight aromatic compounds has attracted much attention^3^. Many bacteria catabolizing lignin-derived aromatic compounds have been identified, and their catabolic enzyme genes have been characterized. These genes are useful for the production of value-added chemicals from heterogeneous lignin-derived aromatic compounds^4^. For example, *Sphingobium* sp. strain SYK-6, an alphaproteobacterium, is a model strain capable of catabolizing various lignin-derived aromatic dimers and monomers and produces 2-pyrone-4,6-dicarboxylic acid (PDC), a platform chemical for the synthesis of functional polymers as an intermediate^5,6^. A significant portion of the SYK-6 genes involved in the catabolism of lignin-derived aromatic compounds have been characterized, and PDC production systems using these genes have been developed^4,7^. Recently, transporter genes for lignin-derived aromatic compounds have also been studied to improve their uptake capacity^8,9^. However, uptake systems, especially uptake across the outer membrane, remain largely unexplored.

The outer membrane transport by Gram-negative bacteria is mediated by porins, substrate-specific channels, and TonB-dependent transporters (TBDTs)^10^. Porins and substrate-specific channels are responsible for nonspecific and specific passive transport, respectively. TBDTs are active transporters that specifically transport relatively large compounds, such as siderophores and vitamin B12, using the energy derived from proton motive force transmitted from the TonB−ExbB−ExbD complex (Ton complex) localized in the inner membrane^11−13^. Since most bacteria have only a few *tonB*-like genes compared to the number of TBDT-like genes, one TonB likely interacts with multiple TBDTs to transfer the energy required for substrate uptake^14,15^. In addition to the above substrates, the involvement of TBDTs in the uptake of oligopeptides, saccharides, and metal ions has been demonstrated, thus, expanding the range of substrates transported by TBDTs^16−18^. It is also known that *tolA, tolQ*, and *tolR*, homologs of *tonB, exbB*, and *exbD*, respectively, constitute the Tol-Pal system, stabilizing the outer membrane structure and is involved in the septum formation during cell division^19^. The introduction of *tolQ* and *tolR* into *exbB* and *exbD* mutants of *Escherichia coli* restored the ability of these mutants to take up group B colicin, indicating that *tolQ* and *tolR* can partially complement the functions of *exbB* and *exbD*, respectively^20^.

The outer membrane transport of aromatic compounds used as carbon and energy sources is mediated by passive transporters, such as the vanillate specific-channel OpdK of *Pseudomonas aeruginosa* PAO1 and naphthalene porin OmpW of *Pseudomonas fluorescens*^21,22^. However, SYK-6 has no OpdK-like genes and only one gene that shows similarity with the known aromatic compound porin (*ompW*)^21,23^, but instead has 74 TBDT-like genes^24^. Recently, we have demonstrated that a TBDT (DdvT) mediates the uptake of 5,5′-dehydrodivanillate (DDVA), a lignin-derived 5,5-type dimer, across the outer membrane in SYK-6. When *ddvT* was overexpressed in an SYK-6 mutant of the PDC hydrolase gene, the PDC production rate improved. Therefore, overexpression of TBDT genes seems effective in improving the efficiency of microbial conversion. Additionally, the Ton complex composed of TonB1−ExbB1−ExbD1−ExbD2 was suggested to be involved in the uptake of DDVA mediated by DdvT. The phylogenetic analysis of 74 TBDT-like genes in SYK-6 with known TBDT genes revealed that 53 TBDT-like genes form two phylogenetically distinct clades from known TBDT genes. Among them, the expression of 12 TBDT genes in SYK-6 was specifically induced by lignin-derived aromatic compounds.

These facts suggest that the Ton system, consisting of TBDTs and Ton complex, plays a vital role in the uptake of lignin-derived aromatic compounds by SYK-6 across the outer membrane. Additionally, the genomes of strains of Sphingomonadaceae, which include many strains capable of degrading recalcitrant aromatic compounds, contain many TBDT-like genes, suggesting that the Ton system is involved in the outer membrane transport of these aromatic compounds^25^. A recent study strongly suggests that TBDTs are involved in the uptake of aromatic compounds, such as benzo[a]pyrene in *Novosphingobium pentaromativorans* US6-1, which degrades polyaromatic hydrocarbons^26^.

This study investigated whether the Ton system is involved in the outer membrane transport of lignin-derived aromatic compounds in SYK-6 by analyzing disruption and overexpression strains of genes presumed to encode components of the Ton complex. Additionally, we predicted the Tol-Pal system genes, which contributed to the stabilization of the outer membrane and characterized their mutants.

## Results and discussion

### The *ompW*-like gene is not required for the conversion of lignin-derived aromatic compounds

SLG_38320 (*ompW*) of SYK-6 shows similarity with known aromatic compound porins (*ompW* family proteins; 30% identity with P0A915 of *Escherichia coli*)^21,23,24^. In our previous study, *ompW* mutant (Δ*ompW*) exhibited comparable growth to the wild type in Wx medium containing 5 mM DDVA, ferulate (FA), vanillin (VN), vanillate (VA), syringaldehyde (SN), syringate (SA), or protocatechuate (PCA)^24^. Here to clarify whether *ompW* is involved in the uptake of lignin-derived aromatic compounds, we used the resting cells of Δ*ompW* to measure their capacity to convert lignin-derived 5,5-, β-O-4-, β-β-, β-5-, and β-1-type dimers [DDVA, guaiacylglycerol-β-guaiacyl ether (GGE), pinoresinol (PR), dehydrodiconiferyl alcohol (DCA), and 1,2-bis(4-hydroxy-3-methoxyphenyl)-propane-1,3-diol (HMPPD), respectively] and monomers [acetovanillone (AV), FA, VN, VA, SN, SA, and PCA] (Fig. S1, S2). No reduction in the conversion capacity of Δ*ompW* compared to the wild type was observed, indicating that *ompW* is unnecessary to uptake lignin-derived aromatic compounds tested in this study.

### The growth of *tonB* mutants on lignin-derived aromatic compounds and their conversion capacity

SYK-6 has six *tonB*-like genes (SLG_14430 [*tonB1*], SLG_34540 [*tonB2*], SLG_36940 [*tonB3*], SLG_37490 [*tonB4*], SLG_01650 [*tonB5*], and SLG_14690 [*tonB6*]), three *exbB*-like genes (SLG_14440 [*exbB1*], SLG_02500 [*exbB2*/*tolQ*], and SLG_10800 [*exbB3*]), and three *exbD*-like genes (SLG_14450 [*exbD1*], SLG_14460 [*exbD2*], and SLG_02490 [*exbD3*/*tolR*]) (Table 1, Fig. S3)^24^. Our previous mutant analysis showed that *tonB2*−*6* were not involved in the uptake of DDVA^24^. *tonB1* that constitutes the *tonB1* operon (*tonB1*−*exbB1*−*exbD1*−*exbD2*) could not be disrupted, implying that *tonB1* is essential for the growth of SYK-6^24,27^. *tonB2* and *fiuA* encoding a TBDT located immediately downstream of *tonB2* play a major role in iron acquisition across the outer membrane of SYK-6^27^. Therefore, the growth capacity of *tonB2* mutant (Δ*tonB2*) was reduced on various carbon sources^24,27^.

**Table 1.** Candidate genes encoding the Ton complex components in *Sphingobium* sp. strain SYK-6.

| Locus tag | Putative function | Most similar protein<sup>a</sup> | Annotation | Strain | Sequence identity (%) | E-value |
| --- | --- | --- | --- | --- | --- | --- |
| **TonB** |  |  |  |  |  |  |
| SLG_01650 (*tonB5*) | TonB | — | — | — | — | — |
| SLG_14430 (*tonB1*) | TonB | — | — | — | — | — |
| SLG_14690 (*tonB6*) | TonB | — | — | — | — | — |
| SLG_34540 (*tonB2*) | TonB | — | — | — | — | — |
| SLG_36940 (*tonB3*) | TonB | — | — | — | — | — |
| SLG_37490 (*tonB4*) | TonB | — | — | — | — | — |
| **ExbB** |  |  |  |  |  |  |
| SLG_02500 (*exbB2/tolQ*) | ExbB/TolQ | P50598 | Tol-Pal system protein TolQ | *Pseudomonas aeruginosa* PAO1 | 41 | 4E-46 |
| SLG_10800 (*exbB3*) | ExbB | — | — | — | — | — |
| SLG_14440 (*exbB1*) | ExbB | P0C7L9 | Biopolymer transport protein ExbB | *Xanthomonas campestris* pv. *campestris* str. ATCC33913 | 38 | 4E-42 |
| **ExbD** |  |  |  |  |  |  |
| SLG_02490 (*exbD3/tolR*) | ExbD/TolR | P0ABV2 | Biopolymer transport protein ExbD | *Escherichia coli* K-12 | 41 | 7E-27 |
| SLG_14450 (*exbD1*) | ExbD | P0C7I9 | Biopolymer transport protein ExbD2 | *Xanthomonas campestris* pv. *campestris* str. ATCC33913 | 29 | 6E-17 |
| SLG_14460 (*exbD2*) | ExbD | P0C7M0 | Biopolymer transport protein ExbD1 | *Xanthomonas campestris* pv. *campestris* str. ATCC33913 | 37 | 3E-24 |

<sup>a</sup>Most similar proteins were searched in the Swiss-Prot database using the BLAST-P program<sup>40</sup> and not displayed if the E-value is greater than 1e-10.

To examine the involvement of *tonB*s in the uptake of lignin-derived aromatic compounds, we measured the growth of *tonB3 tonB4 tonB5 tonB6* quadruple mutant (Δ*tonB3456*) and its *tonB2* mutant (Δ*tonB23456*) in Wx medium containing 5 mM FA, VN, VA, SA, or PCA, or Wx medium containing SEMP (10 mM sucrose, 10 mM glutamate, 0.13 mM methionine, and 10 mM proline). We also evaluated the capacity of resting cells of Δ*tonB3456* and Δ*tonB23456* to convert GGE, PR, DCA, HMPPD, AV, FA, VN, VA, SN, SA, and PCA. Δ*tonB3456* and Δ*tonB23456* showed no reduction in growth on all carbon sources compared with wild type and Δ*tonB2*, respectively (Fig. S4). Additionally, the conversion capacity of Δ*tonB3456* and Δ*tonB23456* to each compound was not reduced compared to the wild type (Fig. 1). These results suggest that *tonB2*−*6* are not involved in the uptake of these lignin-derived aromatic compounds tested in this study.

**Fig. 1.**
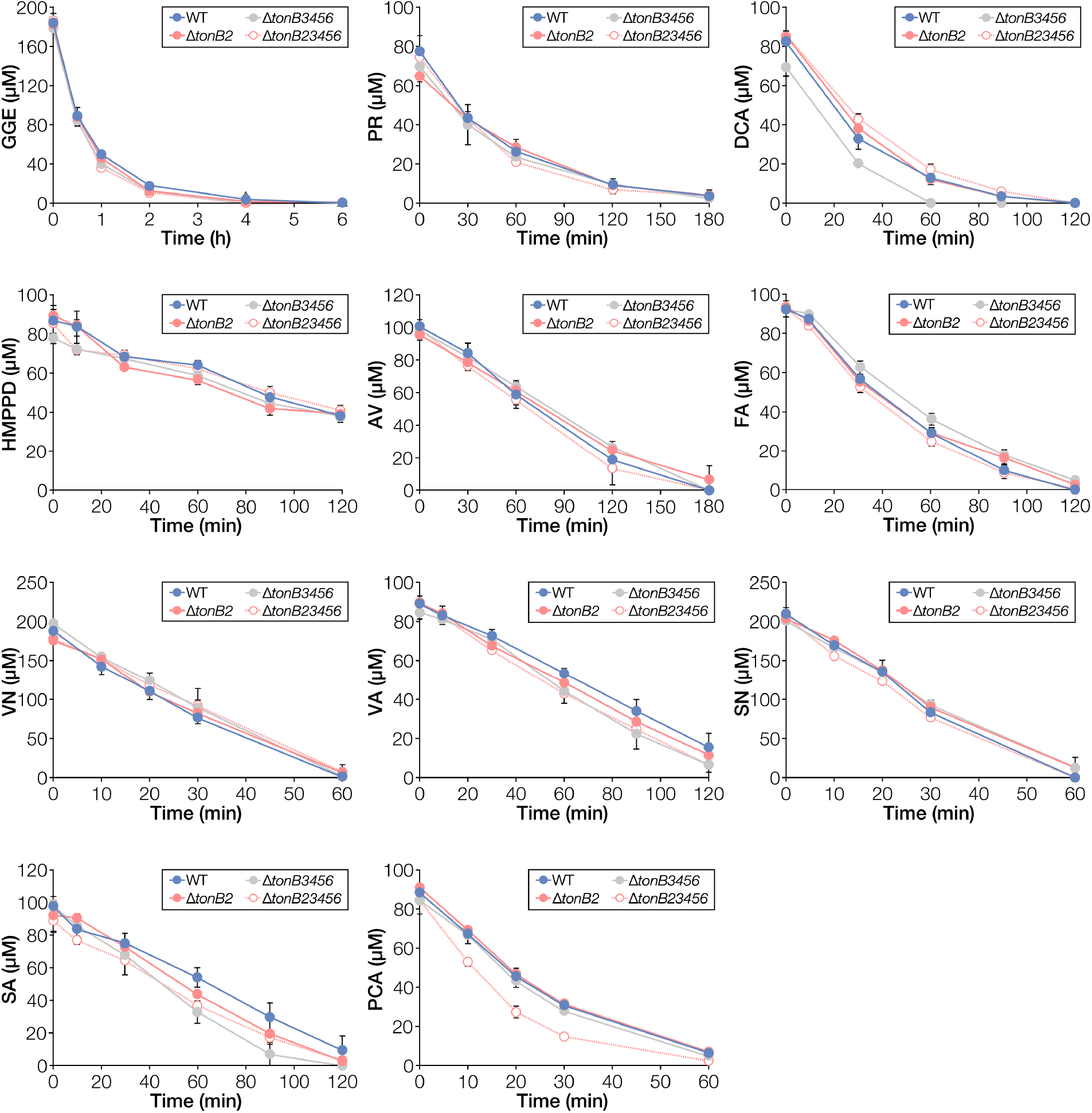
Conversion of lignin-derived aromatic compounds by resting cells of *tonB* multiple mutants. Cells of SYK-6, Δ*tonB2*, Δ*tonB3456*, and Δ*tonB23456* were incubated with 200 µM GGE, 100 µM PR, 100 µM DCA, 100 µM HMPPD, 100 µM AV, 100 µM FA, 200 µM VN, 100 µM VA, 200 µM SN, 100 µM SA, or 100 µM PCA. Portions of the reaction mixtures were collected, and the amount of substrate was measured using HPLC. Each value is the average ± the standard deviation of three independent experiments. GGE, guaiacylglycerol-β-guaiacyl ether; PR, pinoresinol; DCA, dehydrodiconiferyl alcohol; HMPPD, 1,2-bis(4-hydroxy-3-methoxyphenyl)-propane-1,3-diol; AV, acetovanillone; FA, ferulate; VN, vanillin; VA, vanillate; SN, syringaldehyde; SA, syringate; PCA, protocatechuate.

### The growth of *exbB* and *exbD* mutants on lignin-derived aromatic compounds

We tried disrupting three *exbB*-like genes and three *exbD*-like genes and finally obtained *exbB2*/*tolQ, exbB3*, and *exbD3*/*tolR* mutants (Table 1, Fig. S5). *exbB1, exbD1*, and *exbD2*, which form the *tonB1* operon, could not be disrupted, suggesting that this operon plays an essential role in the growth of SYK-6. We measured the growth of *exbB2*/*tolQ* mutant (Δ*exbB2*/*tolQ*), *exbB3* mutant (Δ*exbB3*), and *exbD3*/*tolR* mutant (Δ*exbD3*/*tolR*) in Wx medium containing FA, VN, VA, SA, PCA, or SEMP or diluted LB (Fig. 2). Δ*exbB3* exhibited the same level of growth as the wild type on all substrates. In contrast, the growth of Δ*exbB2*/*tolQ* and Δ*exbD3*/*tolR* was significantly reduced or defective in SEMP, diluted LB, and lignin-derived aromatic compounds except for FA. The growth was restored when the plasmids carrying *exbB2*/*tolQ* and *exbD3*/*tolR* were introduced into Δ*exbB2*/*tolQ* and Δ*exbD3*/*tolR*, respectively (Fig. S6, S7). Therefore, growth retardation or defect was due to the disruption of *exbB2*/*tolQ* and *exbD3*/*tolR*. However, the introduction of *exbB1* into Δ*exbB2*/*tolQ* and *exbD1*-*exbD2* into Δ*exbD3*/*tolR* did not restore their growth (Fig. S6, S7). These results indicate that *exbB1* and *exbD1*/*exbD2* cannot replace the functions of *exbB2*/*tolQ* and *exbD3*/*tolR*, respectively.

**Fig. 2.**
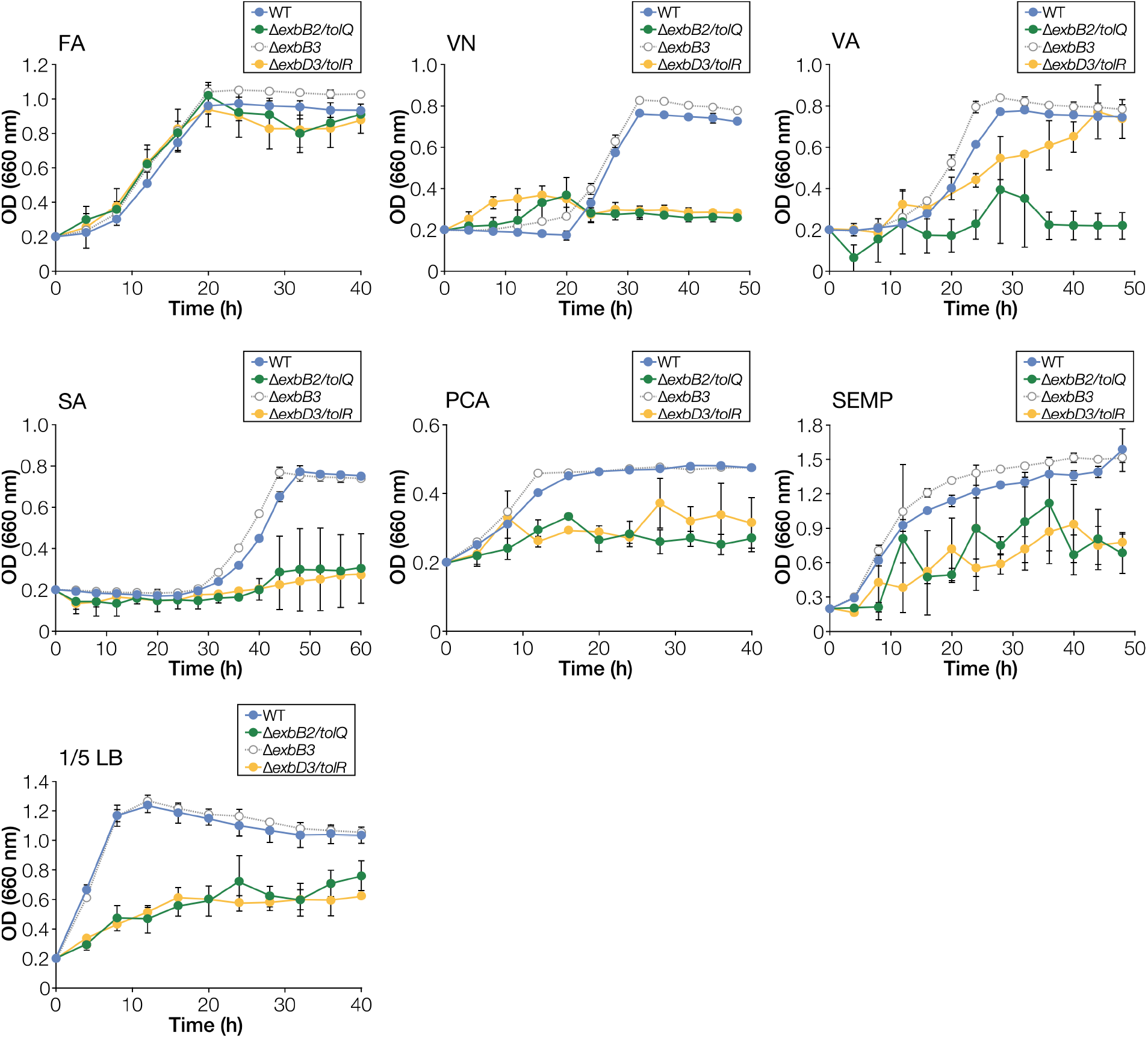
Growth of *exbB* and *exbD* mutants on lignin-derived aromatic compounds. Cells of SYK-6, Δ*exbB2*/*tolQ*, Δ*exbB3*, and Δ*exbD3*/*tolR* were cultured in Wx medium containing 5 mM FA, VN, VA, SA, or PCA, Wx medium containing SEMP, and diluted LB (1/5 LB). Cell growth was monitored by measuring the OD_660_. Each value is the average ± the standard deviation of three independent experiments.

*exbB2*/*tolQ* and *exbD3*/*tolR* were located just upstream of *tolA, tolB*, and *pal*-like genes that constitute the Tol-Pal system (Fig. S3), which plays essential roles in the stabilization of the outer membrane structure and septum formation during cell division. Additionary, *ybgC*, whose relevance to the Tol-Pal system is unknown, was found immediately upstream of *exbB2*/*tolQ*^19,28^. Since a similar set of genes (*ybgC*−*tolQ*−*tolR*−*tolA*−*tolB*−*pal*) has been found in many bacteria as the genes encoding the Tol-Pal system^29^, *exbB2*/*tolQ* and *exbD3*/*tolR* are likely components of the Tol-Pal system. Therefore, disruption of these genes is presumed to have caused instability of the outer membrane and associated growth retardation and defects on various carbon sources. The details of the characterization of Δ*exbB2*/*tolQ* and Δ*exbD3*/*tolR* will be described later.

In ExbB and TolQ of *E. coli*, Thr-148 and Thr-181 (ExbB numbering) are essential for proton translocation and Glu-176 (ExbB numbering) whose function is unclear but essential, are conserved^30,31^. These amino acid residues were conserved in ExbB1 and ExbB2/TolQ of SYK-6 (Fig. S8A). However, Glu-176 is not conserved in ExbB3 of SYK-6. With the fact that the growth capacity of Δ*exbB3* was not reduced on each carbon source compared to the wild type (Fig. 2), likely, ExbB3 does not function as ExbB. In contrast, Asp-25 (ExbD numbering) essential for acquiring proton motive force by ExbD and TolR in *E. coli* was conserved among ExbD1, ExbD2, and ExbD3/TolR of SYK-6 (Fig. S8B).

### The effect of overexpression of the *tonB1* operon genes on the conversion of lignin-derived aromatic compounds

To examine the involvement of the *tonB1* operon genes (*tonB1*−*exbB1*−*exbD1*−*exbD2*) in the uptake of lignin-derived aromatic compounds, each plasmid carrying *tonB1* (pJB-tonB1), *tonB1*−*exbB1*−*exbD1* (pJB-t1-D1), and *tonB1*−*exbB1*−*exbD1*−*exbD2* (pJB-t1-D12) was introduced into SYK-6 cells. We measured the capacity of resting cells of SYK-6 harboring pJB-tonB1, pJB-t1-D1, or pJB-t1-D12 to convert DDVA, GGE, PR, DCA, HMPPD, AV, FA, VN, VA, SN, SA, and PCA. The introduction of pJB-tonB1 increased only the conversion rate of DDVA by ca. 1.8-fold, while the introduction of pJB-t1-D1 or pJB-t1-D12 increased the conversion rate of DDVA, GGE, PR, DCA, HMPPD, FA, VA, SA, and PCA by ca. 1.1−2.6-fold (Table 2, Fig. S9). However, the introduction of these plasmids did not promote the growth of SYK-6 on DDVA, FA, VN, VA, PCA, and SEMP (Fig. S10). On the other hand, the conversion rates of AV, VN, and SN by SYK-6 cells harboring pJB-tonB1, pJB-t1-D1, or pJB-t1-D12 were comparable to those of the control strain. AV, VN, and SN are presumed to be taken up by the outer membrane transporters other than TBDT or diffused without transporters.

**Table 2.** The conversion rates of lignin-derived aromatic compounds by SYK-6 cells overexpressing the *tonB1* operon genes.

| Substrate | WT + pJB861 | WT + pJB-tonB1 | WT + pJB-t1-D1 | WT + pJB-t1-D12 |
| --- | --- | --- | --- | --- |
| DDVA | 7.0 ± 1.1 | 12 ± 0.6** [1.8] | 15 ± 0.1*** [2.2] | 18 ± 1.4*** [2.6] |
| GGE | 45 ± 2.4 | 49 ± 1.7 | 53 ± 1.1* [1.2] | 53 ± 0.2** [1.2] |
| PR | 35 ± 1.2 | 36 ± 0.8 | 43 ± 0.7** [1.2] | 43 ± 2.2* [1.2] |
| DCA | 133 ± 3.4 | 142 ± 7.4 | 164 ± 1.8*** [1.2] | 161 ± 0.5*** [1.2] |
| HMPPD | 98 ± 4.6 | 101 ± 5.8 | 81 ± 13 | 111 ± 4.2* [1.1] |
| AV | 42 ± 4.4 | 41 ± 1.1 | 42 ± 2.6 | 41 ± 1.0 |
| FA | 73 ± 2.3 | 74 ± 1.3 | 74 ± 2.8 | 86 ± 1.0** [1.2] |
| VN | 229 ± 13 | 257 ± 15 | 187 ± 35 | 229 ± 8.0 |
| VA | 38 ± 2.9 | 38 ± 1.9 | 42 ± 1.3 | 47 ± 0.5* [1.2] |
| SN | 210 ± 16 | 197 ± 20 | 212 ± 16 | 193 ± 8.6 |
| SA | 44 ± 4.0 | 44 ± 2.0 | 51 ± 1.7 | 59 ± 1.8** [1.4] |
| PCA | 73 ± 1.8 | 72 ± 2.9 | 75 ± 1.3 | 85 ± 2.2** [1.2] |

To determine whether the enhanced conversion capacity of the strains that overexpress the *tonB1* operon genes was due to their increased uptake capacity, we assessed the DDVA uptake of these strains using a previously constructed DDVA biosensor^9,24^. Consequently, overexpression of *tonB1, tonB1*−*exbB1*−*exbD1*, and *tonB1*−*exbB1*−*exbD1*−*exbD2* increased the DDVA uptake capacity ca. 1.4-, 1.3-, and 1.6-fold, respectively, compared to the control strain (Fig. 3). These results strongly suggest that the *tonB1* operon genes are involved in the uptake of DDVA. Similarly, strains overexpressing the *tonB1* operon genes likely enhanced the conversion of GGE, PR, DCA, HMPPD, FA, VA, SA, and PCA due to the enhanced capacity to take up each substrate. Therefore, these lignin-derived aromatic compounds seem to be taken up by TBDTs.

**Fig. 3.**
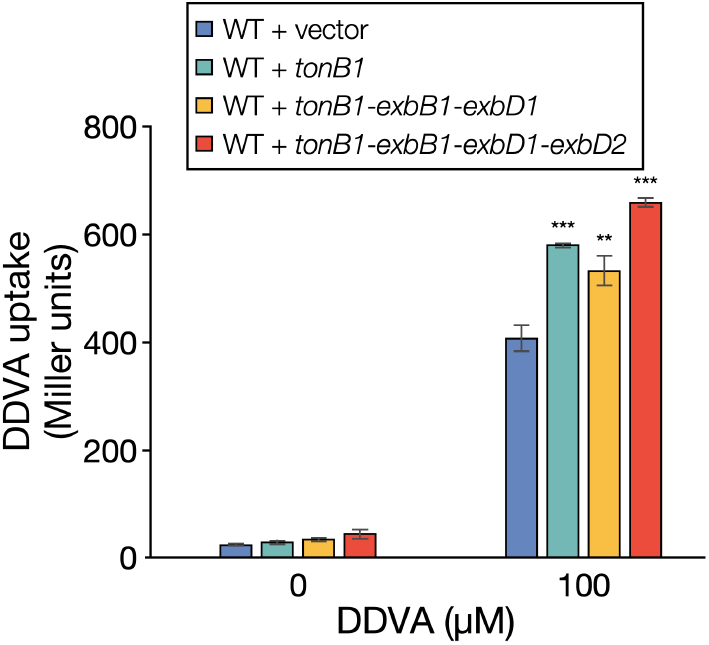
DDVA uptake by SYK-6 cells overexpressing the *tonB1* operon genes. The uptake of DDVA by SYK-6 cells overexpressing the *tonB1* operon genes was evaluated using the DDVA sensor plasmid pS-XR, which was constructed by applying the regulatory system of the DDVA catabolic genes^24^. The β-galactosidase activities of cells of SYK-6(pS-XR + pSEVA338 [vector]), SYK-6(pS-XR + pS-tonB1), SYK-6(pS-XR + pS-t1-D1), and SYK-6(pS-XR + pS-t1-D12) incubated in Wx-SEMP with or without 100 µM DDVA were measured. Each value is the average ± the standard deviation of three independent experiments. Asterisks show statistically significant differences between cells overexpressing the *tonB1* operon genes and vector control cells incubated in the presence of 100 µM DDVA. ^**^, *P* < 0.01, ^***^, *P* < 0.001 (unpaired, two-tailed *t*-test).

We investigated the cellular localization of TonB1. Western blot analysis using anti-TonB1 antibodies against a cell extract and a total membrane fraction obtained from SYK-6 cells grown in LB showed a clear signal in a total membrane fraction, indicating that TonB1 is localized in the cell membrane (Fig. S11).

Sphingomonadaceae, to which SYK-6 belongs, contains many strains capable of degrading recalcitrant aromatic compounds such as lignin-derived aromatic compounds and polycyclic aromatic hydrocarbons^25^. These strains have as many or more TBDT and Ton complex genes as SYK-6^25,27^. Therefore, we examined whether the *tonB1* operon genes of SYK-6 are conserved in the Sphingomonadaceae strains listed in Table S1. These strains have genes showing 43−58% amino acid sequence identity with *tonB1*, 35−81% with *exbB1*, 35−81% with *exbD1*, and 55−75% with *exbD2*. Additionally, these genes have a similar gene organization as the SYK-6 *tonB1* operon (Fig. S12). Based on these facts, it is highly likely that the *tonB1* operon-like genes play a central role in TonB-dependent uptake in these Sphingomonadaceae.

### ExbB2/TolQ and ExbD3/TolR constitute the Tol-Pal system

Since mutations in genes constituting the Tol-Pal system have been reported to cause reduced resistance to detergents^28^, we compared the growth of wild type, Δ*exbB2*/*tolQ*, and Δ*exbD3*/*tolR* in LB with or without 0.3 mg/ml sodium deoxycholate (NaDOC) or 0.05 mg/ml sodium dodecyl sulfate (SDS) (Fig. 4). The addition of NaDOC or SDS to the wild type had little effect on growth; however, the growth of Δ*exbB2*/*tolQ* and Δ*exbD3*/*tolR* was significantly reduced in the presence of the detergents.

**Fig. 4.**
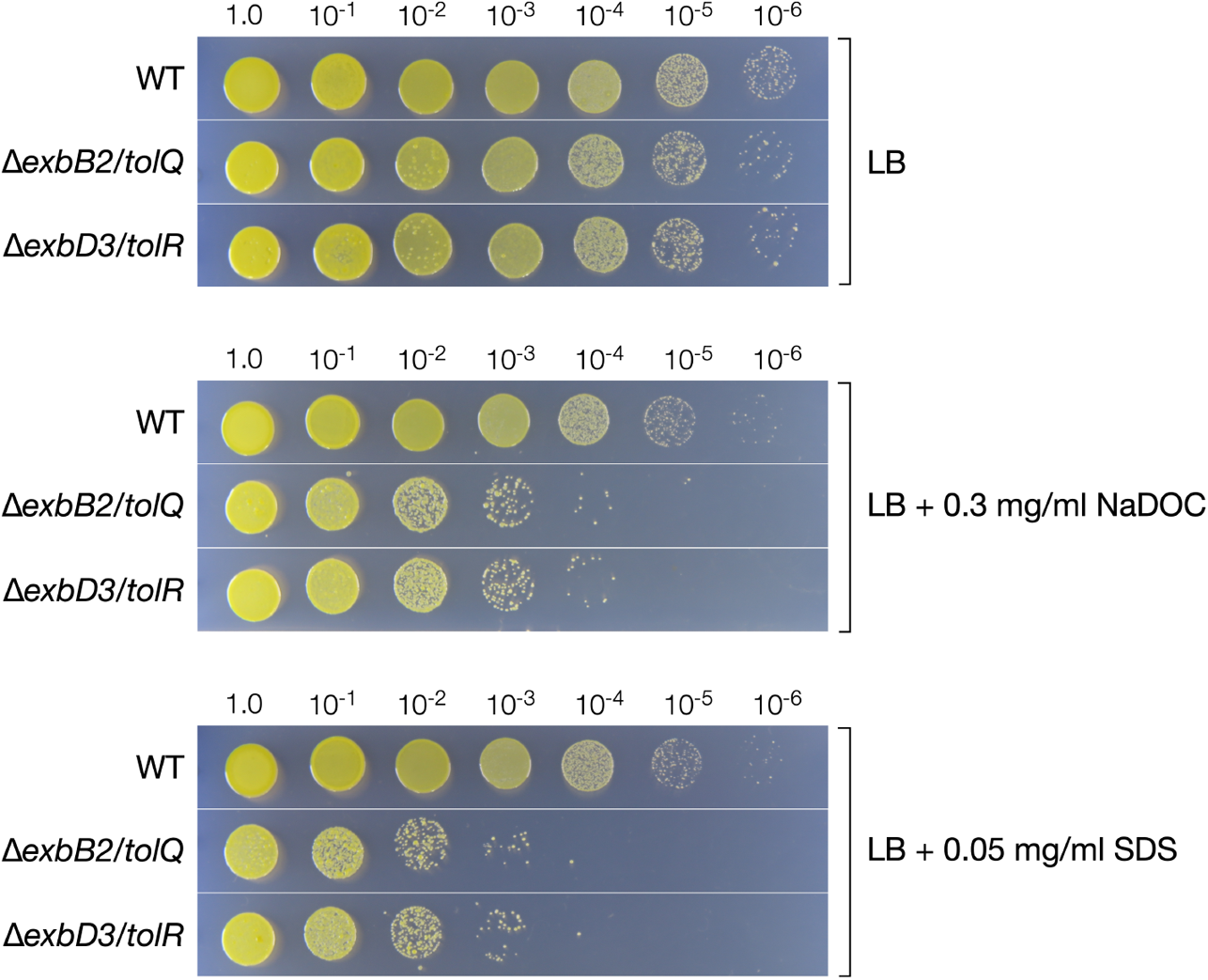
Δ*exbB2*/*tolQ* and Δ*exbD3*/*tolR* cells show susceptibility to detergents. The cells of SYK-6, Δ*exbB2*/*tolQ*, and Δ*exbD3*/*tolR* were grown in LB with or without 0.3 mg ml^−1^ of NaDOC or 0.05 mg ml^−1^ SDS. The cell diluted with Wx buffer (10 µl) were dropped onto LB agar medium and incubated at 30 °C for 72 h.

Mutations in the Tol-Pal system genes cause instability of the outer membrane structure, resulting in the secretion of the outer membrane vesicles (OMVs)^32^. Therefore, we observed the cell surface structure of the wild type, Δ*exbB2*/*tolQ*, and Δ*exbD3*/*tolR* grown in LB using field emission scanning electron microscopy (FE-SEM) (Fig. 5A). The formation of blebs was observed on the cell surface of Δ*exbB2*/*tolQ* and Δ*exbD3*/*tolR*. To examine whether OMVs were secreted into the culture medium of Δ*exbB2*/*tolQ* and Δ*exbD3*/*tolR*, we performed western blot analysis using anti-DdvT antibodies against the culture supernatants of wild type, Δ*exbB2*/*tolQ*, and Δ*exbD3*/*tolR* grown in LB since OMVs are thought to contain outer membrane proteins (Fig. 6, S13)^24,32^. DdvT was detected in the culture supernatants of Δ*exbB2*/*tolQ* and Δ*exbD3*/*tolR*, but not in the culture supernatant of the wild type. Furthermore, DdvT was not detected in the ultracentrifugated supernatant obtained from the cultures of Δ*exbB2*/*tolQ* and Δ*exbD3*/*tolR*, suggesting the presence of outer membrane fractions in their culture supernatants. Western blot analysis using anti-TonB1 antibodies as an inner membrane marker protein showed that TonB1 was detected only in cell extracts of all strains. These results indicate that OMVs are present in the culture supernatants of Δ*exbB2*/*tolQ* and Δ*exbD3*/*tolR*.

**Fig. 5.**
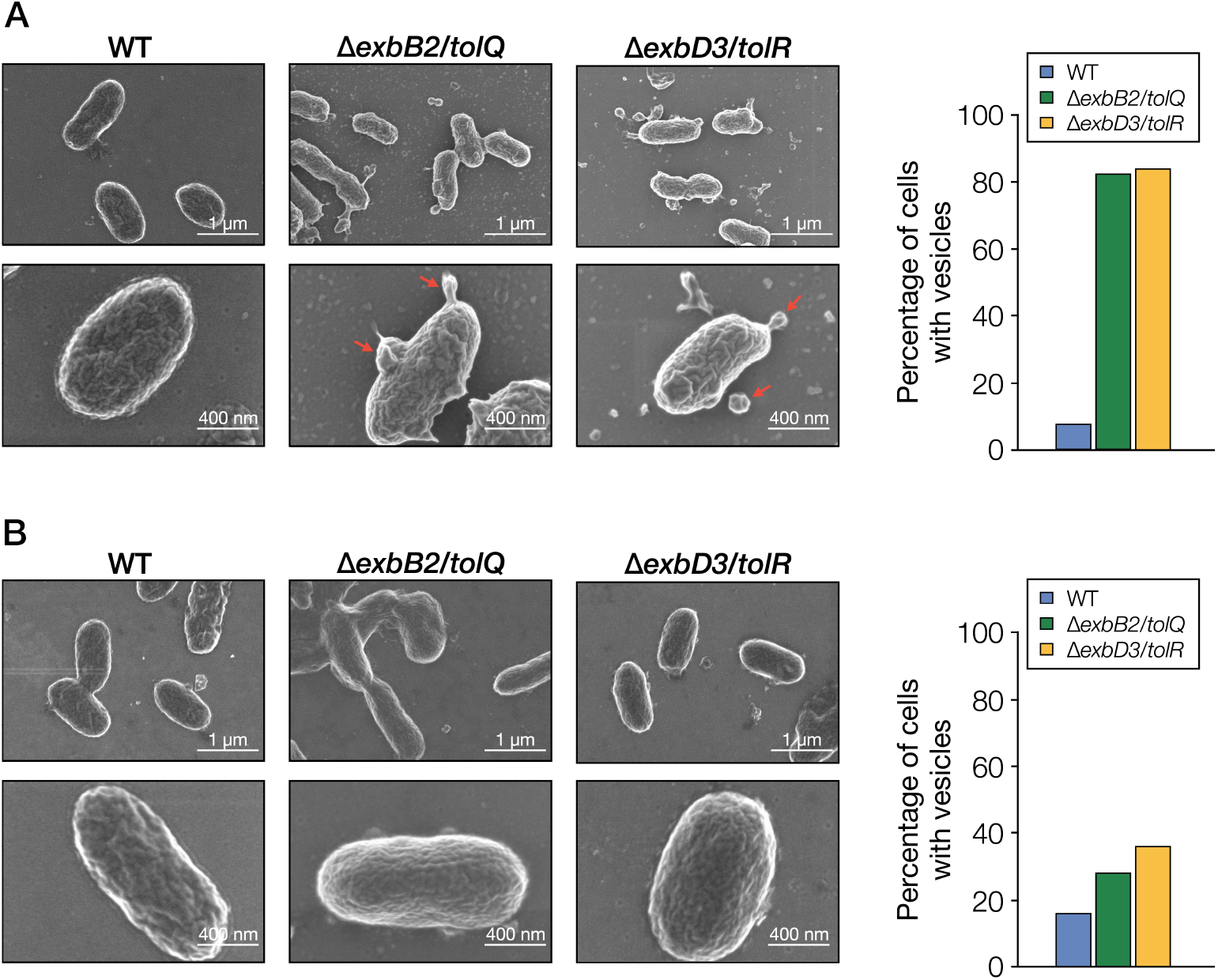
SEM analysis of SYK-6, Δ*exbB2*/*tolQ*, and Δ*exbD3*/*tolR*. (A) FE-SEM images of SYK-6, Δ*exbB2*/*tolQ*, and Δ*exbD3*/*tolR* cells grown in LB. Red arrows indicate blebbing vesicles. (B) FE-SEM images of SYK-6, Δ*exbB2*/*tolQ*, and Δ*exbD3*/*tolR* cells grown in LB with 5 mM FA. The right figure shows the percentage of cells with vesicles. Outer membrane vesiculation was quantified by imaging 50 bacterial cells and expressed as the percentage of cells with OMV.

**Fig. 6.**
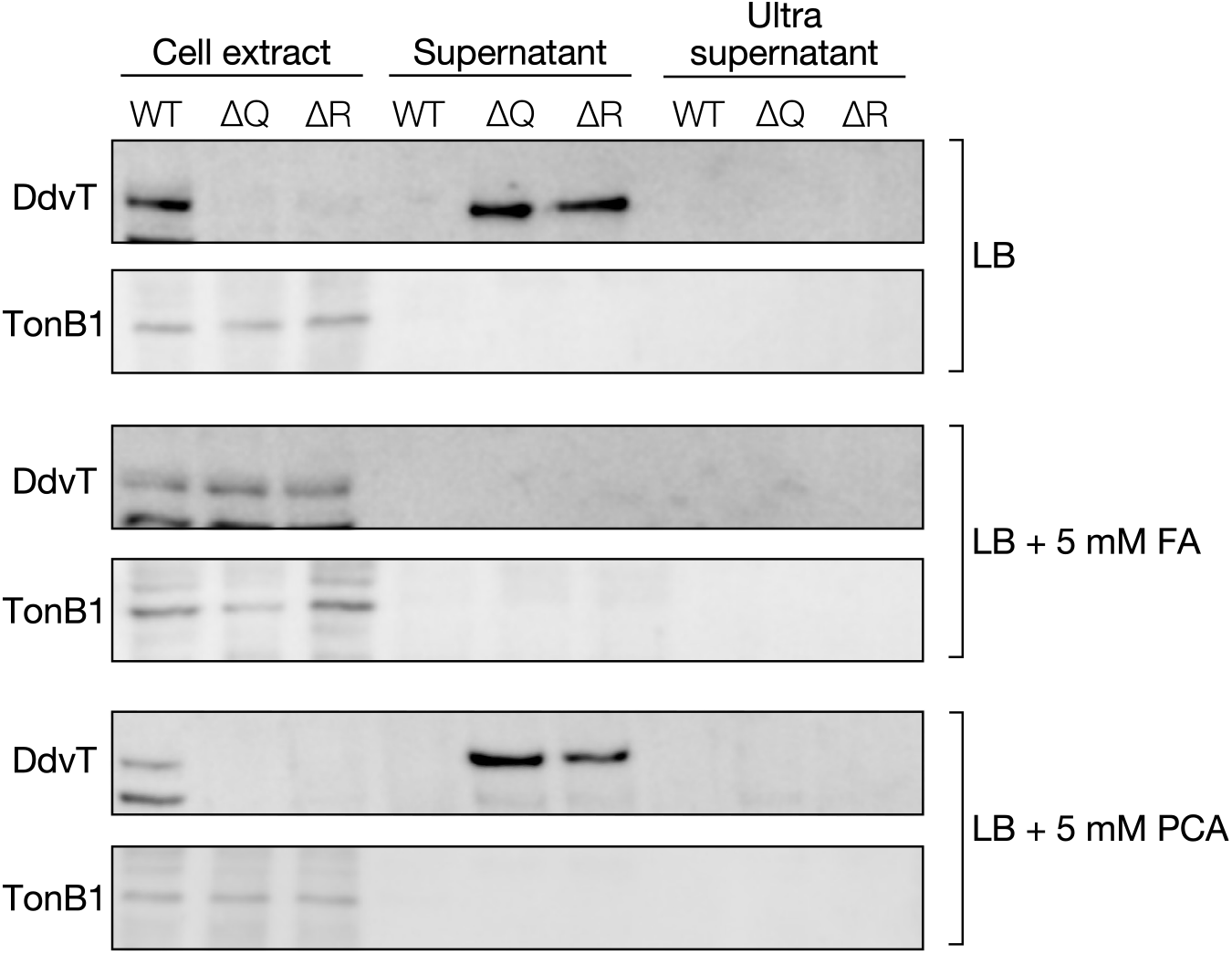
Detection of the outer membrane fraction in the culture supernatants of *exbB2*/*tolQ* and *exbD3*/*tolR* mutants. Cells of the wild type (WT), Δ*exbB2*/*tolQ* (ΔQ), and Δ*exbD3*/*tolR* (ΔR) were grown in LB with or without 5 mM FA or PCA. As shown in Fig. S13, after centrifugation, the supernatants of the cultures were collected (supernatant), and the cell extracts were prepared. The supernatants were further ultracentrifuged, and the resulting supernatants were collected (ultrasupernatant). Western blot analysis using anti-DdvT and anti-TonB1 antibodies was performed against cell extracts (10 μg of protein), 20 µl of the supernatant, and 20 µl of the ultrasupernatant.

FA was the only carbon source that did not reduce the growth of Δ*exbB2*/*tolQ* and Δ*exbD3*/*tolR* (Fig. 2). Thus, vesiculation of these mutants during FA catabolism was evaluated. Interestingly, DdvT was not detected in the culture supernatants of Δ*exbB2*/*tolQ* and Δ*exbD3*/*tolR* grown in LB containing 5 mM FA (Fig. 6). When Δ*exbB2*/*tolQ* and Δ*exbD3*/*tolR* were grown in LB containing 5 mM PCA, a catabolite of FA (Fig. S14A), DdvT was detected in their culture supernatants, suggesting that the outer membrane structure of Δ*exbB2*/*tolQ* and Δ*exbD3*/*tolR* was specifically stabilized during FA catabolism. The observation of the cell surface structure of Δ*exbB2*/*tolQ* and Δ*exbD3*/*tolR* grown in LB containing 5 mM FA showed that the formation of blebs was significantly suppressed during FA catabolism (Fig. 5B). In SYK-6 cells, FA is converted to VN through the addition of coenzyme A (CoA) to the Cγ position and the subsequent release of acetyl-CoA (Fig. S14A)^33^. To get insight into why the growth capacity of Δ*exbB2*/*tolQ* and Δ*exbD3*/*tolR* was not reduced when FA was used as a carbon source (Fig. 2), we constructed a double mutant of *exbD3*/*tolR* and the PDC hydrolase gene *ligI* (Δ*exbD3*/*tolR ligI*) (Fig. S5, S14A). This strain produces acetyl-CoA during FA catabolism, but the conversion of VN from FA stops at the PDC. Growth measurement of Δ*exbD3*/*tolR ligI* in the presence or absence of FA showed that the addition of more than 1 mM FA compensated for the growth in diluted LB (Fig. S14B). In contrast, the addition of VA and PCA did not positively affect the growth of Δ*exbD3*/*tolR ligI* in diluted LB. Based on these results, the continuous supply of acetyl-CoA produced in the second step of FA catabolism to fatty acid biosynthesis may reduce the damage to the outer membrane caused by the disruption of *exbB2*/*tolQ* and *exbD3*/*tolR* (Fig. S14A). However, further investigation is needed to elucidate the actual mechanism.

In the Sphingomonadaceae strains that can degrade aromatic compounds listed in Table S2, genes with high similarity amino acid sequences (*exbB2*/*tolQ*, 56−63% identity; *exbD3*/*tolR*, 46−62% identity; *tolA*, 33−52% identity; *tolB*, 58−73% identity; *pal*, 57−70% identity) and gene organization with the Tol-Pal system genes of SYK-6 were conserved (Fig. S15). This high degree of conservation suggests that these genes play similar roles to the corresponding SYK-6 genes in these bacterial strains.

## Conclusions

Overexpression of the *tonB1* operon genes in SYK-6 enhanced the capacity to convert at least five lignin-derived dimers and four monomers. These results strongly suggest that TBDTs mediate the uptake of these lignin-derived aromatic compounds, and the Ton complex composed of TonB1−ExbB1−ExbD1−ExbD2 is involved in their uptake (Fig. 7). Besides, *exbB2*/*tolQ* and *exbD3*/*tolR* constitute the Tol-Pal system stabilizing the outer membrane structure. Based on this study’s results, it is expected that microbial conversion systems with enhanced uptake capacity will be developed through the identification of TBDT genes involved in the outer membrane transport of each lignin-derived aromatic compound.

**Fig. 7.**
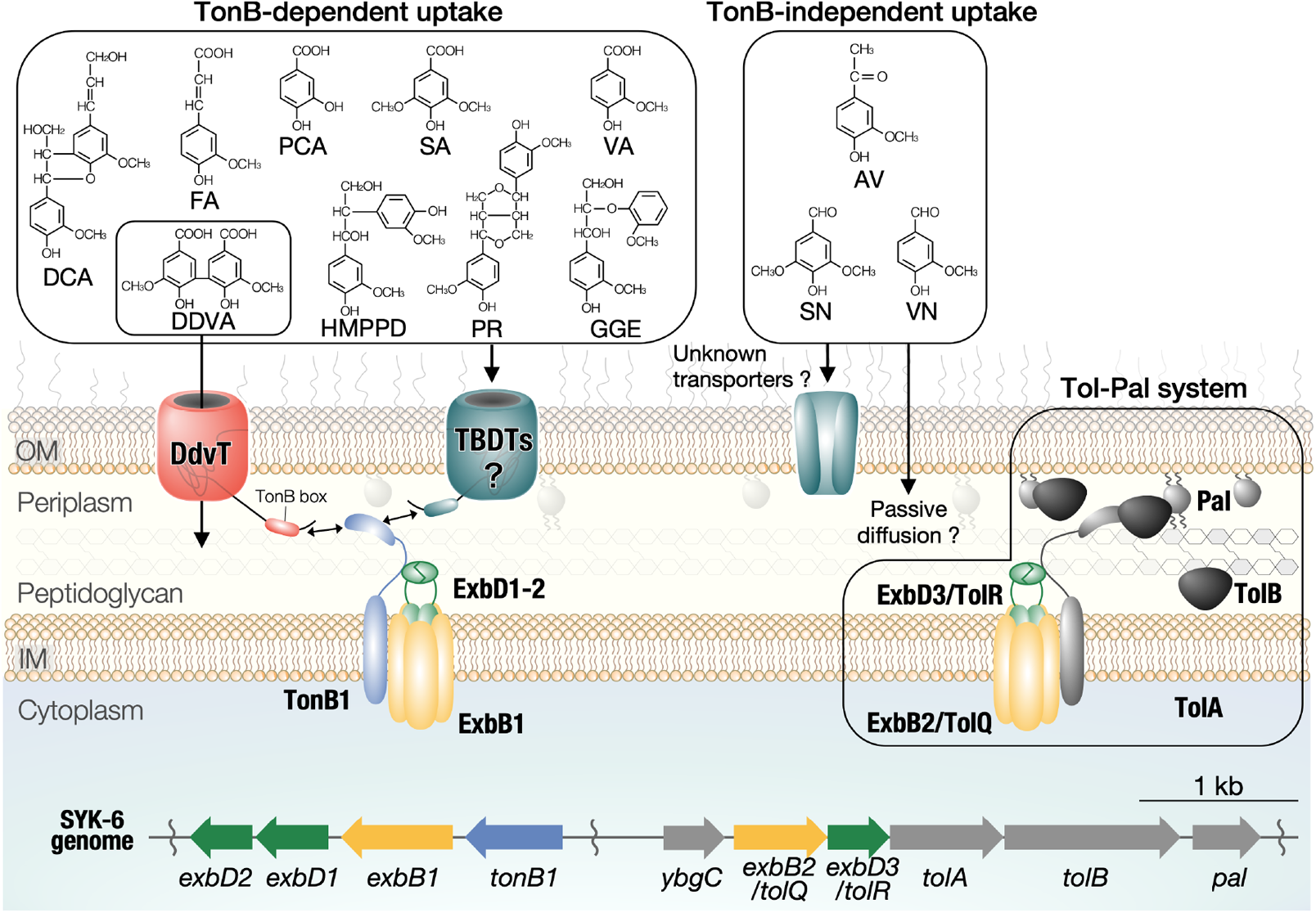
Proposed outer membrane transport of lignin-derived aromatic compounds in SYK-6. Outer membrane transport of GGE, PR, DCA, HMPPD, FA, VA, SA, and PCA is thought to be mediated by TBDTs, and the involvement of TonB1−ExbB1−ExbD1−ExbD2 as the Ton complex is strongly suggested. AV, VN, and SN are presumed to be taken up by transporters other than TBDT or passive diffusion. ExbB2/TolQ and ExbD3/TolR are the components of the Tol-Pal system that stabilizes the outer membrane structure. OM, outer membrane; IM, inner membrane.

## Methods

### Bacterial strains, plasmids, culture conditions, and substrates

The strains and plasmids used in this study are listed in Table S3. *Sphingobium* sp. SYK-6 (NBRC 103272/JCM 17495) and its mutants were grown at 30 °C with shaking (160 rpm) in LB or Wx minimal medium with SEMP^34^. Media for SYK-6 transformants and mutants were supplemented with 50 mg l^−1^ kanamycin (Km) or 30 mg l^−1^ chloramphenicol (Cm). *E. coli* strains were cultured in LB at 37 °C. Media for *E. coli* transformants carrying antibiotic resistance markers were supplemented with 25 mg l^−1^ Km or 30 mg l^−1^ Cm. Lignin-derived compounds were purchased from Sigma-Aldrich and Tokyo Chemical Industry Co., Ltd. DDVA, PR, DCA, and HMPPD were synthesized as previously described^35−38^.

### Mutant construction

Plasmids for gene disruption were constructed by amplifying ca. 1-kb fragments carrying upstream and downstream regions of each gene by PCR with SYK-6 genome DNA as a template and the primer pairs (Table S4). The resultant PCR fragments were inserted into the BamHI site in pAK405 by NEBuilder HiFi DNA assembly cloning kit (New England Biolabs, Inc.). These plasmids were independently introduced into SYK-6 and its mutant cells by triparental mating, and candidate mutants were isolated as previously described^39^. The disruption of the genes was confirmed by colony PCR using primer pairs (Table S4). The plasmids for gene complementation and overexpression (Table S3) were introduced into SYK-6 and the mutant cells by electroporation.

### Sequence analysis

DNA sequencing of PCR amplification products was conducted by Eurofins Genomics. Sequence analysis was conducted using the MacVector program v.15.5.2. Additionally, sequence similarity searches, multiple alignments, and pairwise alignments were conducted using the BLAST^40^, Clustal Omega^41^, and EMBOSS programs^42^, respectively.

### Growth measurement

Cells of SYK-6, its mutants, and complemented strains were grown in LB for 24 h. The cells were harvested by centrifugation at 4,800 × *g* for 5 min, washed twice with Wx buffer, and resuspended in 3 ml of the same buffer. The cells were then inoculated into diluted LB, Wx medium containing SEMP, or Wx medium containing 5 mM DDVA, FA, VN, VA, SA, or PCA to an OD_660_ of 0.2. Since SYK-6 exhibits auxotrophy for methionine when grown in a methoxy-group-free substrate, 0.13 mM methionine was added to the medium to grow on PCA. Cells were incubated at 30 °C with shaking (60 rpm), and cell growth was monitored every hour by measuring the OD_660_ with a TVS062CA biophotorecorder (Advantec Co., Ltd.). For the assay of complemented strains, Km and 1 mM *m*-toluate (an inducer of the P_*m*_ promoter in pJB861) were added to the medium; for the assay of SYK-6 cells overexpressing the *tonB1* operon genes, Km and 0.5 mM *m*-toluate were added to the medium.

### Resting cell assay

Cells of SYK-6 and its mutant were grown in LB for 20 h and harvested by centrifugation at 14,100 × *g* for 5 min. The cells were washed twice with 50 mM Tris-HCl buffer (pH 7.5) and resuspended in 1 ml of the same buffer. For conversion of AV, cells were grown in Wx-SEMP until OD_600_ reached 0.5 and then incubated in the presence of 5 mM AV for 12 h. The cells were then incubated in 50 mM Tris-HCl buffer (pH 7.5) with 100 μM DDVA (OD_600_ of the cells, 5.0; reaction time, 6 h), 200 µM GGE (5.0; 6 h), 100 µM PR (0.5; 3 h), 100 µM DCA (0.5; 2 h), 100 μM HMPPD (2.0; 2 h), 100 µM AV (0.5; 3 h), 100 µM FA (1.0; 2 h), 200 µM VN (0.2; 1 h), 100 µM VA (2.0: 2 h), 200 µM SN (0.4; 1 h), 100 µM SA (2.0; 2 h), or 100 µM PCA (2.0; 1 h). Samples were collected periodically, and the reactions were stopped by centrifugation at 18,800 × *g* for 10 min. The supernatants were diluted 5-fold in water, filtered, and analyzed using high-performance liquid chromatography (HPLC). For the analysis of SYK-6 cells harboring plasmids carrying the *tonB1* operon genes, the cells grown in LB containing Km and 0.5 mM *m*-toluate were employed. These cells were incubated with lignin-derived aromatic compounds under the conditions described above, with the following exceptions. For the conversion of 100 µM DDVA, 100 µM GGE, 100 µM PR, and 100 µM PCA, cells with OD_600_ of 5.0, 3.0, 0.2, and 1.0, respectively, were used and incubated with each substrate for 5, 3, 4, and 2 h.

### HPLC conditions

HPLC analysis was conducted using an Acquity UPLC system (Waters Corporation) with a TSKgel ODS-140HTP column (2.1 by 100 mm; Tosoh Corporation). All analyses were conducted at a flow rate of 0.5 ml min^−1^. The mobile phase was a mixture of solution A (acetonitrile containing 0.1% formic acid) and solution B (water containing 0.1% formic acid) under the following conditions: i) conversion of DDVA, 0−2.5 min, 15% A; ii) conversion of GGE, 0−3.2 min, linear gradient from 5 to 40% A; 3.2−6.0 min, decreasing gradient from 40 to 5% A; 6.0−7.0 min, 5% A. iii) conversion of PR, 0−5.0 min, 20%A. iv) conversion of DCA, 0−3.0 min, 25% A. v) conversion of HMPPD, 0−2.5 min, 10% A. vi) conversion of AV, 0−2.0 min, 15% A. vii) conversion of FA, 0−2.5 min, 15% A. viii) conversion of VN and SN, 0−2.0 min, 15% A. ix) conversion of VA and SA, 0−2.0 min, 10% A. x) conversion of PCA, 0−1.5 min, 10% A. DDVA, GGE, PR, DCA, HMPPD, AV, FA, VN, VA, SN, SA, and PCA were detected at 265, 270, 280, 277, 278, 276, 322, 280, 260, 308, 275, and 260 nm, respectively.

### DDVA uptake assay

Cells of SYK-6 harboring pS-XR and pS-tonB1, pS-t1-D1, or pS-t1-D12 grown in LB containing Km and Cm for 20 h were harvested by centrifugation at 4,800 × *g* for 5 min, washed twice with Wx buffer, and resuspended in 1 ml Wx-SEMP. The cells were then inoculated into Wx-SEMP containing 0.5 mM *m*-toluate with or without 100 µM DDVA to an OD_600_ of 2.0. Samples were incubated at 30 °C with shaking (1,500 rpm) for 3 h. The β-galactosidase activity of the cells was measured using 2-nitrophenyl-β-D-galactopyranoside as the substrate, according to a modified Miller assay [https://openwetware.org/wiki/Beta-Galactosidase_Assay_(A_better_Miller)]^9^. β-galactosidase activity was expressed as Miller units.

### Detergent resistance assay

Cells of SYK-6 and its mutants were grown in LB for 20 h were harvested by centrifugation at 4,800 × *g* for 5 min and resuspended in Wx buffer. The cells were inoculated into LB with or without 0.05 mg ml^−1^ SDS or 0.3 mg ml^−1^ NaDOC to an OD_600_ of 0.2 and incubated at 30 °C for 24 h. The cultured cells were then serially diluted with Wx buffer, and 10 µl of each cell suspension was dropped onto an LB plate. The plates were incubated at 30 °C for 72 h.

### Western blot analysis

A peptide corresponding to residues 199–210 (QAGNPIRTKDRR) of TonB1 was synthesized and used as an antigen to obtain antisera for TonB1 in rabbits (Cosmo Bio, Inc.). Anti-TonB1-peptide antibodies were obtained by purifying the antiserum using peptide affinity column chromatography (Cosmo Bio, Inc.). Anti-DdvT antibodies were obtained in a previous study^24^. Total membrane fractions were prepared as described previously from SYK-6 cells incubated in LB for 20 h^24^. TonB1 and DdvT were detected by western blot analysis using anti-TonB1 antibodies (0.11 µg ml^−1^) and anti-DdvT antibodies (0.25 µg ml^−1^) as described previously^24^. Horseradish peroxidase-conjugated goat anti-rabbit IgG antibodies (Invitrogen, 0.2 µg ml^−1^) were used as the secondary antibodies. Protein concentrations were determined by the Bradford method using a Bio-Rad protein assay kit or Lowry’s assay with a DC protein assay kit (Bio-Rad Laboratories). TonB1 and DdvT were detected using the ECL Western Blotting Detection System (GE Healthcare) with a LumiVision PRO image analyzer (Aisin Seiki Co., Ltd).

### Detection of outer membrane vesiculation of Δ*exbB2*/*tolQ* and Δ*exbD3*/*tolR*

Cells of SYK-6 and its mutants were grown in LB with or without 5 mM FA or PCA for 20 h were harvested by centrifugation at 19,000 × *g* for 10 min. The supernatant of the centrifugated culture was filtered with a 0.45 µm membrane filter (supernatant fraction). Then, the supernatant fraction was ultracentrifuged at 120,000 × *g* for 60 min to obtain ultracentrifuged supernatant (ultrasupernatant fraction). The cells were washed twice with 50 mM Tris-HCl buffer (pH 7.5) containing 150 mM NaCl and resuspended in the same buffer containing 1 mM phenylmethylsulfonyl fluoride. The cells were disrupted by sonication, and cell lysate was obtained. Each fraction was separated by SDS-7.5% or 15% polyacrylamide gel electrophoresis and performed western blot analysis using anti-TonB1 antibodies and anti-DdvT antibodies as described above.

### FE-SEM analysis

Cells of SYK-6 and its mutants grown on LB plates with or without 5 mM FA for 48 h were deposited on poly-D-lysine coated glass (Cosmo Bio, Inc.) and fixed with 2.0% glutaraldehyde solution in 200 mM sodium phosphate buffer (pH 7.5) for 2 h at room temperature. Fixed cells were washed twice with the same buffer and dehydrated in an ethanol gradient of 25%, 50%, 75%, 90%, and 100% for 15 min per step. Samples were dried using an evaporator for 8 h at 30 °C and sputter-coated with Os. Cell images were obtained using a HITACHI SU8230 field emission SEM with a beam accelerating voltage of 1.0 kV.

### Statistics and reproducibility

All results were obtained from *n* = 3 independent experiments. Statistical tests were performed using GraphPad Prism9 (GraphPad software). Unpaired, two-tailed *t*-test were used as shown in figure legends. *P* < 0.05 was considered statistically significant.

## Data availability

All data supporting this study are available within the article and its Supplementary Information or are available from the corresponding author upon request.

## Acknowledgments

This work was supported by JSPS KAKENHI Grant Numbers 15H04473 (E.M.), 19H02867 (E.M.), 19J11312 (M.F.), and 21J00894 (M.F.). Part of this work resulted from using research equipment shared in MEXT Project for promoting public utilization of advanced research infrastructure (Program for the SHARE, GIGAKU-Innovation Equipment Sharing Network) at Nagaoka University of Technology, Grant Number [JPMXS0430300121].

## Author contributions

E.M. supervised the project. M.F., N.K., and E.M. designed the study and wrote the manuscript. M.F. performed the experiments with the following exceptions. S.Y. and K.S. analyzed PR, DCA, and AV conversion by *tonB* mutants and overexpression strains of the *tonB1* operon genes. M.K. and M.F. performed FE-SEM analysis. S.H. synthesized DCA, PR, and HMPPD. S.Y. performed a statistical analysis of the conversion of lignin-derived aromatic compounds by overexpression strains of the *tonB1* operon genes. S.Y., K.S., and K.M. helped to interpret the data and discussed the results. All authors read and approved the manuscript.

