## Supplementary information for "Functional roles of multiple Ton complex genes in a *Sphingobium* degrader of lignin-derived aromatic compounds"

**Contents list**

Supplementary tables: Table S1-S4
Supplementary figures: Fig. S1-S15
References

**Table S1. Conservation of the *tonB1* operon genes of *Sphingobium* sp. SYK-6 in selected Sphingomonadaceae strains**

| Species and strain | *tonB1* operon | TonB1 | | ExbB1 | | ExbD1 | | ExbD2 | |
|---|---|---|---|---|---|---|---|---|---|
| | | Accession no.[a] | Sequence identity (%)[b] | Accession no.[a] | Sequence identity (%)[b] | Accession no.[a] | Sequence identity (%)[b] | Accession no.[a] | Sequence identity (%)[b] |
| *Novosphingobium aromaticivorans* DSM12444 | + | ABD24466 | 47 | ABD24465 | 66 | ABD24464 | 43 | ABD24463 | 56 |
| *Novosphingobium pentaromativorans* US6-1 | + | AIT80170 | 46 | AIT80171 | 65 | AIT80172 | 43 | AIT80173 | 55 |
| *Novosphingobium resinovorum* SA1 | + | AOR77983 | 43 | AOR77984 | 64 | AOR77985 | 40 | AOR77986 | 55 |
| *Novosphingobium* sp. PP1Y | + | CCA94311 | 47 | CCA94312 | 65 | CCA94313 | 43 | CCA94314 | 55 |
| *Sphingobium chlorophenolicum* L-1 | + | AEG50516 | 58 | AEG50517 | 80 | AEG50518 | 76 | AEG50519 | 75 |
| *Sphingobium japonicum* UT26S | + | BAI95564 | 58 | BAI95565 | 81 | BAI95566 | 75 | BAI95567 | 74 |
| *Sphingobium* sp. YBL2 | + | AJR24448 | 58 | AJR24449 | 80 | AJR24450 | 73 | AJR24451 | 75 |
| *Sphingomonas wittichii* RW1 | + | ABQ66489 | 44 | ABQ66490 | 69 | ABQ66491 | 67 | ABQ66492 | 65 |
| *Blastomonas natatoria* DSM 3183 | + | PXW69510 | 52 | PXW69511 | 71 | PXW69512 | 55 | PXW69513 | 66 |
| *Novosphingobium nitrogenifigens* DSM 19370 | + | EGD60281 | 43 | EGD60282 | 35 | EGD60284 | 48 | EGD60285 | 60 |
| *Sphingobium yanoikuyae* ATCC 51230 | + | EKU75209 | 59 | EKU75210 | 79 | EKU75211 | 76 | EKU75212 | 72 |
| *Sphingopyxis alaskensis* RB2256 | + | ABF52528 | 52 | ABF52527 | 62 | ABF52526 | 60 | ABF52525 | 68 |

[a]Most similar proteins were searched using the BLAST-P program[1] showing the E-value is greater than 1e-10.

[b]Amino acid sequence identity was calculated by the EMBOSS Needle pairwise alignment program.

**Table S2. Conservation of the Tol-Pal system genes of *Sphingobium* sp. SYK-6 in selected Sphingomonadaceae and other bacterial strains**

| Species and strain | Tol-Pal cluster | ExbB2/TolQ | | ExbD3/TolR | | TolA | | TolB | | Pal | |
|---|---|---|---|---|---|---|---|---|---|---|---|
| | | Accession no. | Sequence identity (%)[a] | Accession no. | Sequence identity (%)[a] | Accession no. | Sequence identity (%)[a] | Accession no. | Sequence identity (%)[a] | Accession no. | Sequence identity (%)[a] |
| *Novosphingobium aromaticivorans* DSM12444 | + | ABD25459 | 58 | ABD25460 | 49 | ABD25461 | 45 | ABD25462 | 61 | ABD25463 | 64 |
| *Novosphingobium pentaromativorans* US6-1 | + | AIT80992 | 56 | AIT80993 | 54 | AIT80994 | 40 | AIT80995 | 58 | AIT80996 | 64 |
| *Novosphingobium resinovorum* SA1 | + | AOR78292 | 56 | AOR78293 | 50 | AOR78294 | 36 | AOR78295 | 59 | AOR78296 | 64 |
| *Novosphingobium* sp. PP1Y | + | CCA91793 | 56 | CCA91794 | 48 | CCA91795 | 37 | CCA91796 | 58 | CCA91797 | 64 |
| *Sphingobium chlorophenolicum* L-1 | + | AEG49724 | 62 | AEG49725 | 59 | AEG49726 | 48 | AEG49727 | 70 | AEG49728 | 70 |
| *Sphingobium japonicum* UT26S | + | BAI98251 | 63 | BAI98252 | 61 | BAI98253 | 47 | BAI98254 | 68 | BAI98255 | 66 |
| *Sphingobium* sp. YBL2 | + | AJR22959 | 63 | AJR22960 | 62 | AJR22961 | 47 | AJR22962 | 70 | AJR22963 | 66 |
| *Sphingomonas wittichii* RW1 | + | ABQ68495 | 59 | ABQ68494 | 54 | ABQ68493 | 37 | ABQ68492 | 59 | ABQ68491 | 63 |
| *Blastomonas natatoria* DSM 3183 | + | PXW74505 | 61 | PXW74504 | 58 | PXW74503 | 40 | PXW74502 | 66 | PXW74501 | 59 |
| *Novosphingobium nitrogenifigens* DSM 19370 | + | EGD57947 | 58 | EGD57948 | 46 | EGD57949 | 38 | EGD57950 | 59 | EGD57951 | 64 |
| *Sphingobium yanoikuyae* ATCC 51230 | + | EKU74714 | 63 | EKU74713 | 61 | EKU74712 | 52 | EKU74711 | 73 | EKU74710 | 64 |
| *Sphingopyxis alaskensis* RB2256 | + | ABF52054 | 59 | ABF52053 | 54 | ABF52052 | 33 | ABF52051 | 63 | ABF52050 | 57 |
| *Escherichia coli* K-12 | + | P0ABU9 | 33 | P0ABV6 | 26 | P19934 | 13 | P0A855 | 27 | P0A912 | 34 |
| *Pseudomonas aeruginosa* PAO1 | + | P50598 | 36 | P50599 | 35 | P50600 | 20 | P50601 | 30 | Q9I4Z4 | 38 |
| *Caulobacter crescentus* NA1000 | + | ACL96805 | 39 | ACL96804 | 39 | ACL96803 | 30 | ACL96802 | 45 | ACL96801 | 44 |

[a]Amino acid sequence identity was calculated by the EMBOSS Needle pairwise alignment program.

### Table S3. Strains and plasmids used in this study

| Strains or plasmids | Relevant characteristic(s)[a] | Reference or source |
|---|---|---|
| Strains | | |
| *Sphingobium* sp. | | |
| SYK-6 | Wild type; Nal$^r$ Sm$^r$ | 2 |
| SME057 | SYK-6 derivative; ΔSLG_38320 (*ompW*); Nal$^r$ Sm$^r$ | 3 |
| SME097 | SYK-6 derivative; ΔSLG_34540 (*tonB2*); Nal$^r$ Sm$^r$ | 3 |
| SME291 | SYK-6 derivative; ΔSLG_10800 (*exbB3*); Nal$^r$ Sm$^r$ | This study |
| SME295 | SYK-6 derivative; ΔSLG_02490 (*exbD3*/*tolR*); Nal$^r$ Sm$^r$ | This study |
| SME296 | SYK-6 derivative; ΔSLG_02500 (*exbB2*/*tolQ*); Nal$^r$ Sm$^r$ | This study |
| SME303 | SYK-6 derivative; Δ*tonB3456*; Nal$^r$ Sm$^r$ | 3 |
| SME304 | SYK-6 derivative; Δ*tonB23456*; Nal$^r$ Sm$^r$ | 3 |
| SME352 | SYK-6 derivative; Δ *exbD3*/*tolR* and *ligI*; Nal$^r$ Sm$^r$ | This study |
| *Escherichia coli* | | |
| HB101 | *recA13 supE44 hsd20 ara-14 proA2 lacY1 galK2 rpsL20 xyl-5 mtl-1* | 4 |
| NEB 10-beta | *araD*139 Δ(*ara-leu*)7697 *fhuA lacX74 galK* (ϕ80 Δ*lacZ* Δ*M15*) *recA1 endA1 nupG rpsL* (Sm$^r$) Δ(*mrr-hsdRMS-mcrBC*) | New England Biolabs |
| Plasmids | | |
| pRK2013 | Tra$^+$ Mob$^+$ ColE1 replicon; Km$^r$ | 5 |
| pJB861 | RK2 ori broad-host-range expression vector; Km$^r$ $P_m$ *xylS* | 6 |
| pAK405 | Plasmid for allelic exchange and markerless gene deletions in Sphingomonads; Km$^r$ | 7 |
| pSEVA225 | RK2 ori *lacZ* promoter probe broad host range vector; Km$^r$ | 8 |
| pSEVA338 | pBBR1 ori broad-host-range expression vector; Cm$^r$ $P_m$ *xylS* | 8 |
| pAK-02490 | pAK405 with a 1.9-kb deletion cassette carrying up- and downstream regions of *tolR*/*exbD3* | This study |
| pAK-02500 | pAK405 with a 2.0-kb deletion cassette carrying up- and downstream regions of *tolQ*/*exbB2* | This study |
| pAK-10800 | pAK405 with a 2.0-kb deletion cassette carrying up- and downstream regions of *exbB3* | This study |
| pAK-ligI | pAK405 with a 2.0-kb deletion cassette carrying up- and downstream regions of *ligI* | 9 |
| pS-XR | pSEVA225 with a 0.8-kb PCR amplicon carrying *ddvR* and *ligXa* promoter regions | 10 |
| pS-tonB1 | pSEVA338 with a 0.7-kb fragment carrying *tonB1* | 3 |
| pJB-tonB1 | pJB861 with a 0.7-kb NotI-SacI fragment carrying *tonB1* from pS-tonB1 | 3 |
| pS-t1-D1 | pSEVA338 with a 2.1-kb fragment carrying *tonB1*, *exbB1*, and *exbD1* | 3 |
| pJB-t1-D1 | pJB861 with a 2.1-kb NotI-SacI fragment carrying *tonB1*, *exbB1*, and *exbD1* from pS-t1-D1 | 3 |
| pS-t1-D12 | pSEVA338 with a 2.6-kb fragment carrying *tonB1*, *exbB1*, *exbD1*, and *exbD2* | 3 |
| pJB-t1-D12 | pJB861 with a 2.6-kb NotI-SacI fragment carrying *tonB1*, *exbB1*, *exbD1*, and *exbD2* from pS-t1-D12 | 3 |
| pJB-exbB1 | pJB861 with a 0.8-kb fragment carrying *exbB1* | This study |
| pJB-exbD12 | pJB861 with a 1.0-kb fragment carrying *exbD1* and *exbD2* | This study |
| pJB-*exbB2*/*tolQ* | pJB861 with a 0.7-kb fragment carrying *exbB2*/*tolQ* | This study |
| pJB-*exbD3*/*tolR* | pJB861 with a 0.5-kb fragment carrying *exbD3*/*tolR* | This study |

[a]Nal$^r$, Sm$^r$, Km$^r$, and Cm$^r$, resistance to nalidixic acid, streptomycin, kanamycin, and chloramphenicol, respectively.

**Table S4. Primers used in this study**

| Target gene | Primer | Sequences (5′ to 3′) |
|---|---|---|
| For gene disruption | | |
| pAK-02490 | Dis_TopF | CGGTACCCGGGGATCGCGTCTCGACATTCTGCT |
| (*exbD3*/*tolR*) | Dis_TopR | GTGTGATGGGCGAACTGA |
| | Dis_BotF | TCAGTTCGCCCATCACACGCGCTGTGACCATGAAGA |
| | Dis_BotR | CGACTCTAGAGGATCGGACGATGACCTGCTCGT |
| pAK-02500 | Dis_TopF | CGGTACCCGGGGATCGCATCACTCTCCAGTCCC |
| (*exbB2*/*tolQ*) | Dis_TopR | GCCGTCATCGCCTACAAC |
| | Dis_BotF | GTTGTAGGCGATGACGGCGCCGAGCATGACGATCTT |
| | Dis_BotR | CGACTCTAGAGGATCTCGCCAGCCCGATGAAAA |
| pAK-10800 | Dis_TopF | CGGTACCCGGGGATCTCGGCGTCTTCAATCCAC |
| (*exbB3*) | Dis_TopR | AGGACCGTCCAGAGAACC |
| | Dis_BotF | GGTTCTCTGGACGGTCCTGGTGGGCGTTGTCGCTTA |
| | Dis_BotR | CGACTCTAGAGGATCGTGGTGGAAAATCGACGG |
| For confirmation of gene disruption | | |
| *exbD3*/*tolR* | exbD3_F | GAGCGCGACGTCGAAAAG |
| *exbB2*/*tolQ* | exbB2_F | TCAGTTCGCCCATCACACGCGCTGTGACCATGAAGA |
| *exbB3* | exbB3_F | AACTGGCCTCGCGCTATG |
| Construction of plasmids | | |
| pJB-exbB1 | Forward | GCCTAGGCCGCGGCCGCGCGCGGCCACTTCACTGACTGG |
| | Reverse | ATCCCCGGGTACCGAGCTCGTCAGGCCTTGGCAGCAGC |
| pJB-exbD12 | Forward | GCCTAGGCCGCGGCCGCGCGGGCTACGGCCCAGAAGTAG |
| | Reverse | TACCGAGCTCGAATTTCAGAACACGCCGCCGTA |
| pJB-exbD3/tolR | Forward | TCACCATGGGAAGCTTCGTGCACCACGTTCAGCCGGGA |
| | Reverse | GAATTCCTGCAGGATATCTGCTATTGATCGCCAGATGAACGCG |
| pJB-exbB2/tolQ | Forward | TCACCATGGGAAGCTTCGTGTCAGACAAGCCCGGAGAG |
| | Reverse | GAATTCCTGCAGGATATCTGTCAGGCTTCAAGCTCCCG |

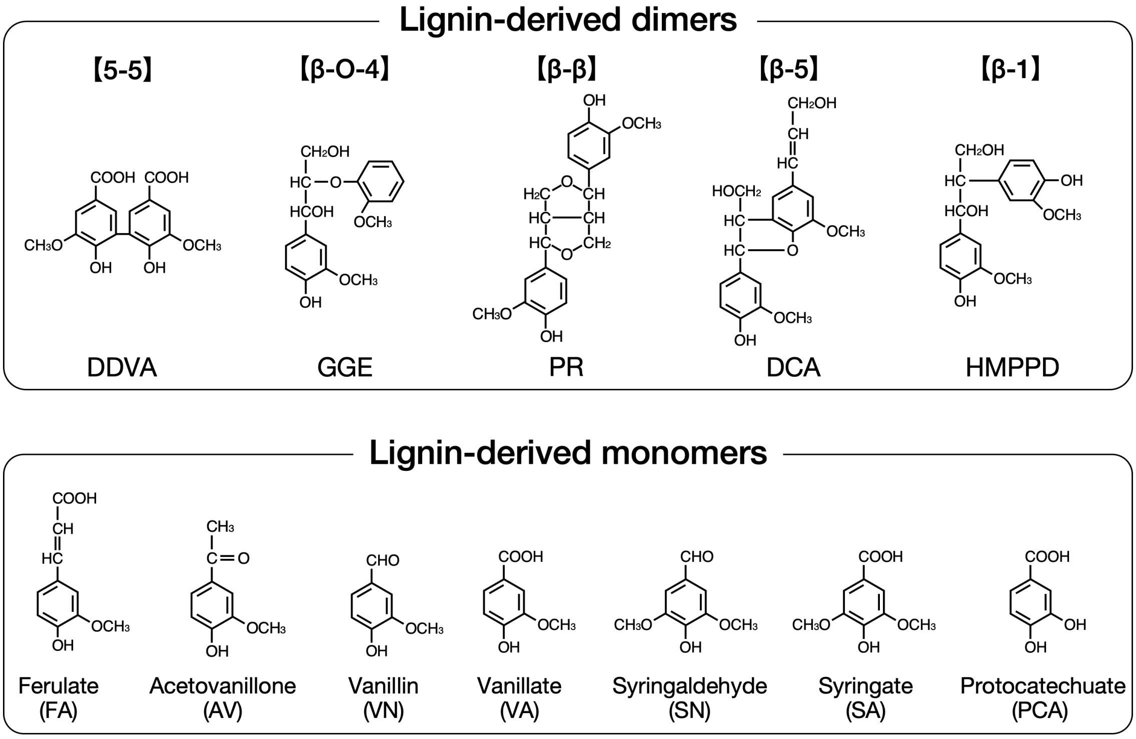


**Fig. S1 Chemical structures of lignin-derived dimers and monomers used in this study.** DDVA, 5,5′-dehydrodivanillate; GGE, guaiacylglycerol-β-guaiacyl ether; PR, pinoresinol; DCA, dehydrodiconiferyl alcohol; HMPPD, 1,2-bis(4-hydroxy-3-methoxyphenyl)-propane-1,3-diol.

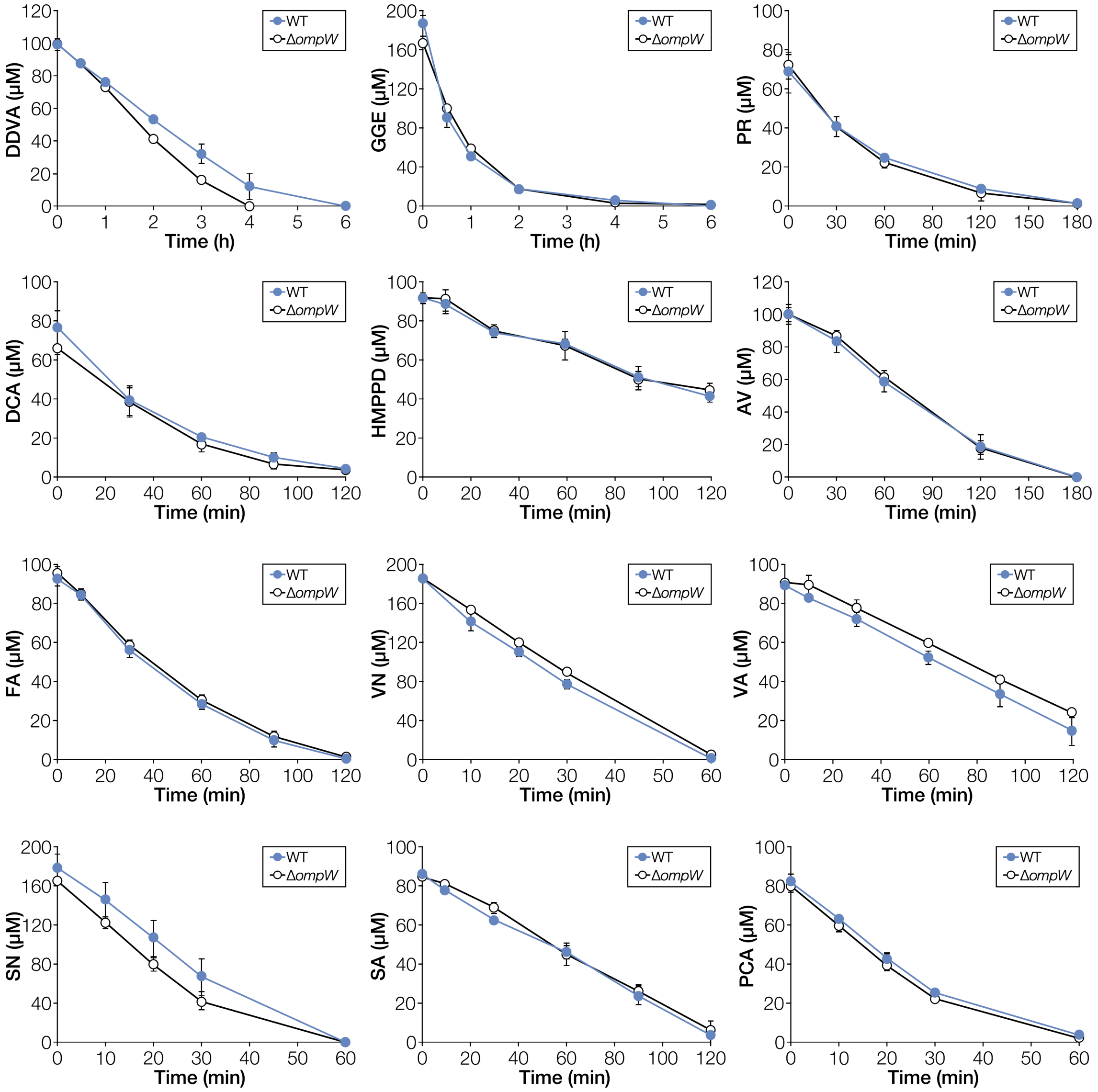


**Fig. S2 Conversion of lignin-derived aromatic compounds by resting cells of Δ*ompW*.** Cells of SYK-6 and Δ*ompW* were incubated with 100 μM DDVA, 200 μM GGE, 100 μM PR, 100 μM DCA, 100 μM HMPPD, 100 μM AV, 100 μM FA, 200 μM VN, 100 μM VA, 200 μM SN, 100 μM SA, and 100 μM PCA, respectively. Portions of the reaction mixtures were collected, and the amount of substrate was measured using HPLC. Each value is the average ± the standard deviation of three independent experiments.

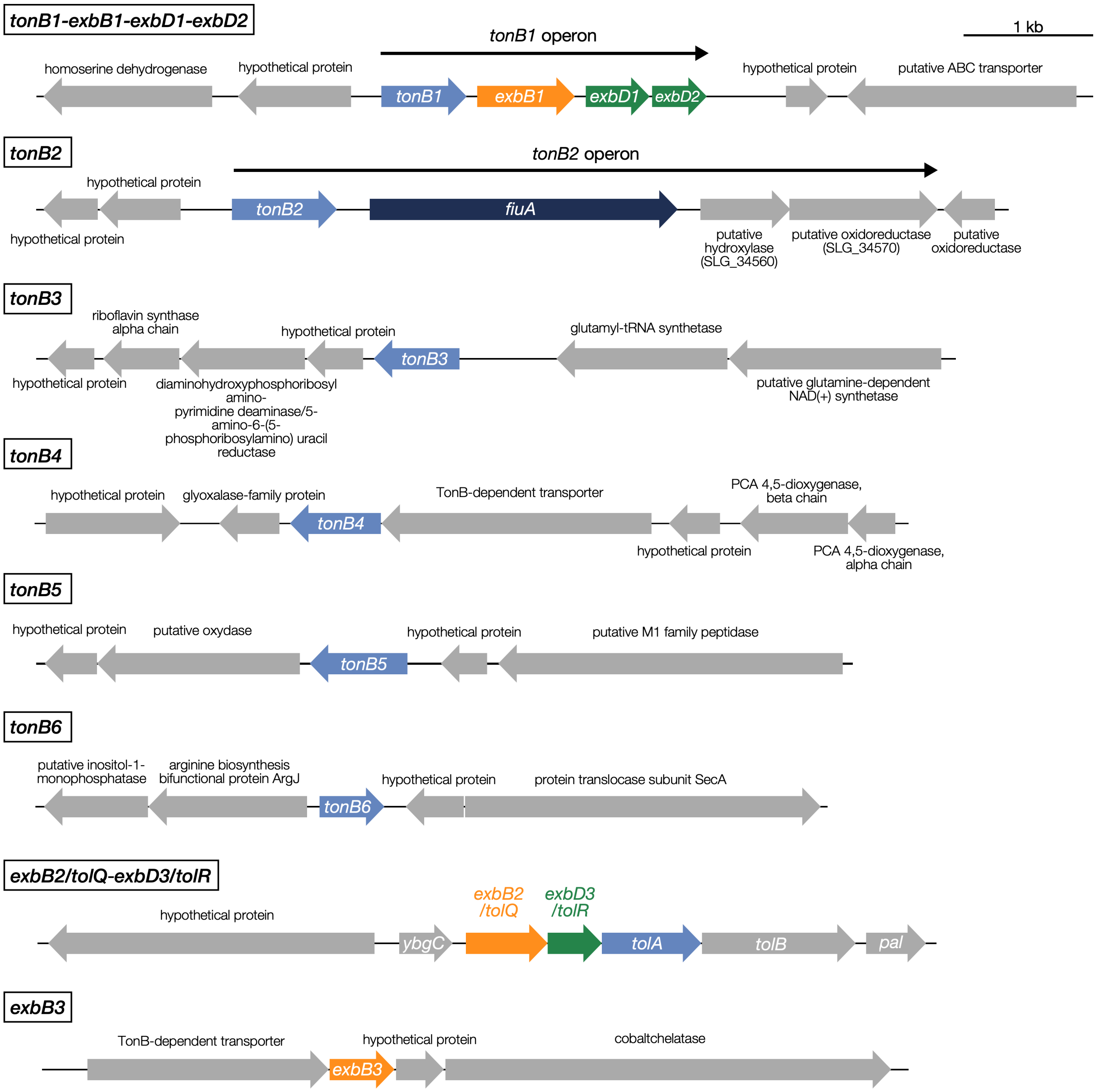


**Fig. S3 Organization of the putative *tonB*, *exbB*, and *exbD* genes in the SYK-6 genome.** *tonB1*–*exbB1*–*exbD1*–*exbD2* and *tonB2*–*fiuA*–SLG_34560–SLG_34570 each form an operon[3,11].

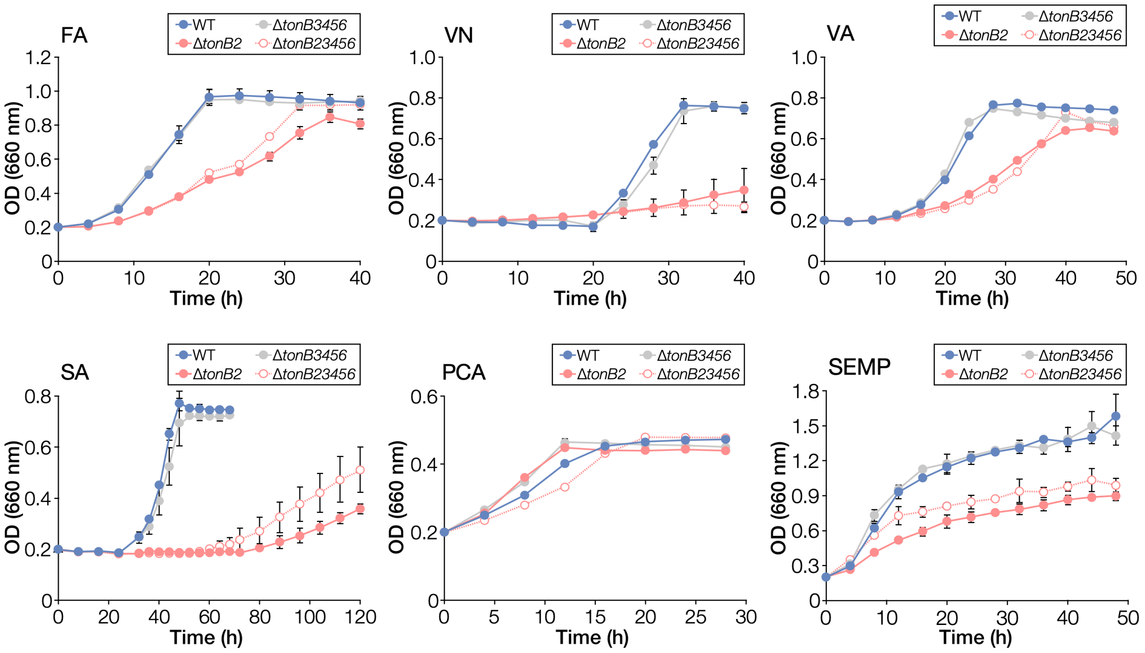


**Fig. S4 Growth of *tonB* multiple mutants on lignin-derived aromatic compounds.** Cells of SYK-6, Δ*tonB2*, Δ*tonB3456*, and Δ*tonB23456* were incubated in Wx medium containing 5 mM FA, VN, VA, SA, or PCA, and Wx medium containing SEMP. Cell growth was monitored by measuring the $OD_{660}$. Each value is the average ± the standard deviation of three independent experiments. The growth data on VA, PCA, and SEMP were reproduced from our previous paper (M. Fujita et al., *Sci. Rep.*, 2020, 10, 12177, https://doi.org/10.1038/s41598-020-68984-2)[11].

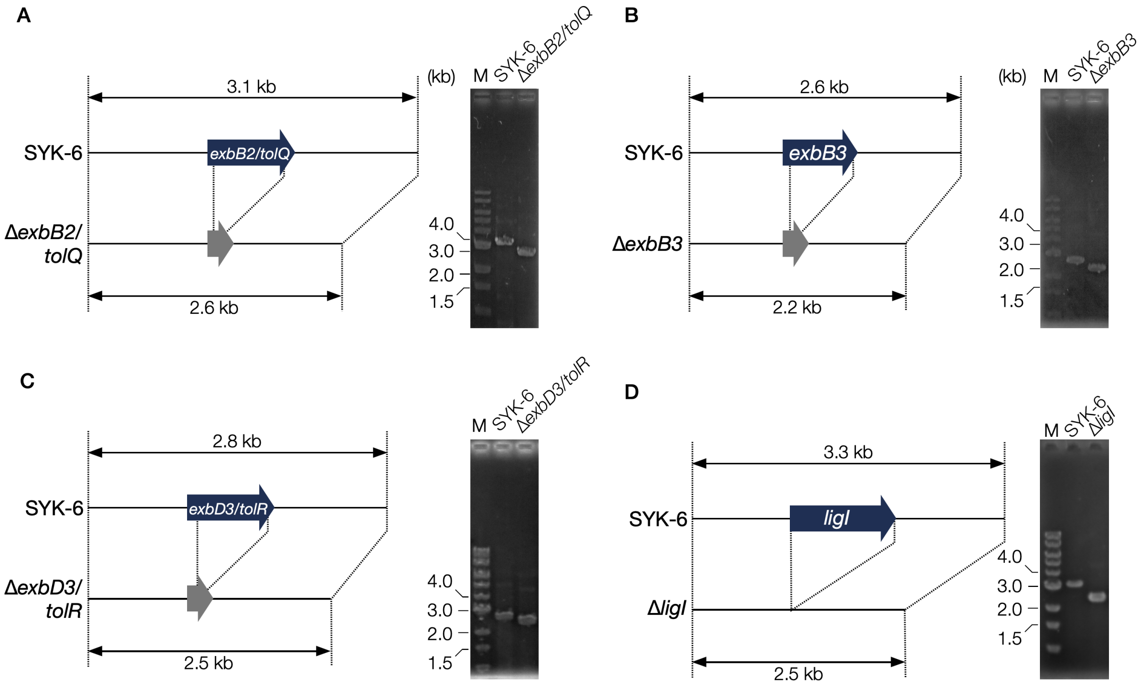


**Fig. S5 Construction of mutants.** (Left) Schematic representations of the disruption of *exbB2/tolQ* (A), *exbB3* (B), *exbD3/tolR* (C), and *ligI* (D). (Right) Colony PCR analysis of each mutant using the primer pairs shown in Table S4. M, molecular size markers.

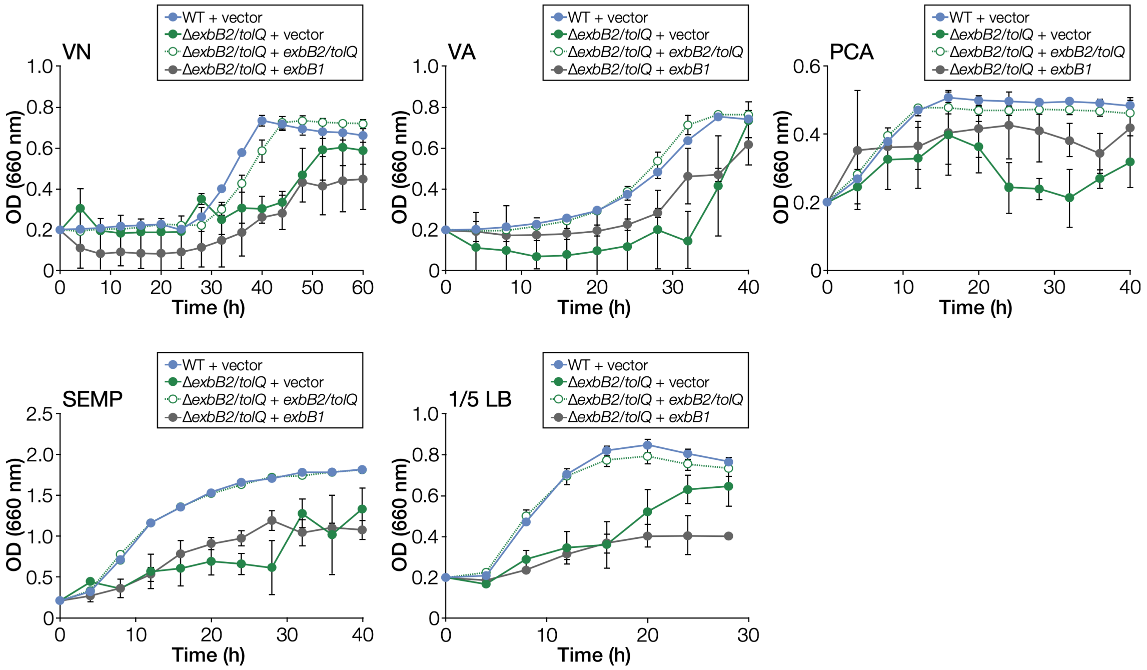


**Fig. S6 Growth complementation of *exbB2*/*tolQ* mutant.** Cells of SYK-6(pJB861), Δ*exbB2*/*tolQ*(pJB861), Δ*exbB2*/*tolQ*(pJB-exbB2/tolQ), and Δ*exbB2*/*tolQ*(pJB-exbB1) were incubated in Wx medium containing 1 mM *m*-toluate and 5 mM VN, VA, or PCA, Wx medium containing 1 mM *m*-toluate and SEMP, and diluted LB containing 1 mM *m*-toluate. Cell growth was monitored by measuring the $OD_{660}$. Each value is the average ± the standard deviation of three independent experiments.

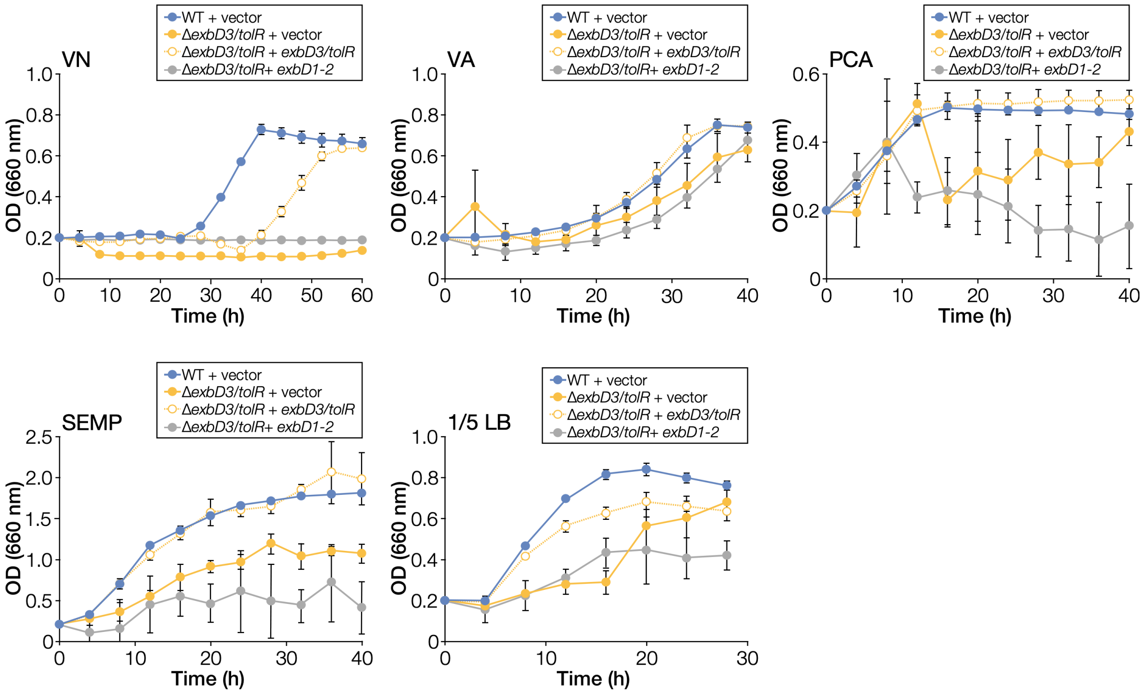


**Fig. S7 Growth complementation of *exbD3*/*tolR* mutant.** Cells of SYK-6(pJB861), Δ*exbD3*/*tolR*(pJB861), Δ*exbD3*/*tolR*(pJB-exbD3/tolR), and Δ*exbD3*/*tolR*(pJB-exbD12) were incubated in Wx medium containing 1 mM *m*-toluate and 5 mM VN, VA, or PCA, Wx medium containing 1 mM *m*-toluate and SEMP, and diluted LB containing 1 mM *m*-toluate. Cell growth was monitored by measuring the $OD_{660}$. Each value is the average ± the standard deviation of three independent experiments.

**A**

```
ExbB1_SYK-6       GLFGTVIGIYRALIKIGASGQASIDAVAGPVGEALIMTALGLAVAVPAVLAYNWLQRRNK 208
ExbB2/TolQ_SYK-6  GLFGTVWGIMRSFTAIAGEQNTSLAVVAPGIAEALFATAIGLFAAIPAVIAYNRLSHGVN 214
ExbB3_SYK-6       GLMGTLIPMATALSGLA---RGDLQILASNMVIAFSSTVVGLAVGVVA---YLVAMVREG 155
ExbB_E. coli      GLFGTVWGIMNSFIGIAQTQTTNLAVVAPGIAEALLATAIGLVAAIPAVVIYNVFARQIG 203
TolQ_E. coli      GLFGTVWGIMHAFIALGAVKQATLQMVAPGIAEALIATAIGLFAAIPAVMAYNRLNQRVN 200
                  **:**:  :  ::  :.      :  :*  :  *:  *.:** ..: *   *
```

**B**

```
ExbD1_SYK-6       ----MAMSAGGGGDDAPMSDINTTPLVDVMLVLLIIFLIAVPVVIQTVEVNLPKIAFEPT 56
ExbD2_SYK-6       -----MAMSGGRDDGEPMMEMNTTPLIDVMLVLLIMFIITIPIQTHAVKIDLPQNADAPQ 55
ExbD3/TolR_SYK-6  MSMSLPPGRRGGGRRAPMAEINVTPLVDVMLVLLIIFMVTAPLLVTGVPIRLPESRARAL 60
ExbD_E. coli      MAMHLNENL---DDNGEMHDINVTPFIDVMLVLLIIFMVAAPLATVDVKVNLPASTSTPQ 57
TolR_E. coli      -----MARARGRGRRDLKSEINIVPLLDVLLVLLLIFMATAPIITQSVEVDLPDATESQA 55
                              .      ::* .*::**:****::*: : *:    * : **
```

**Fig. S8 Conserved amino acid residues between ExbB and TolQ and between ExbD and TolR.** (A) T148, E176, and T181 (ExbB_*E. coli* numbering) are essential for ExbB[12]. (B) D25 (ExbD_*E. coli* numbering) is essential for ExbD[12]. Multiple alignments were constructed using the Clustal Omega program[13]. Accession numbers: ExbB_*E. coli*, P0ABU7; TolQ_*E. coli*, P0ABU9; ExbD_*E. coli*, P0ABV2; TolR_*E. coli*, P0ABV6.

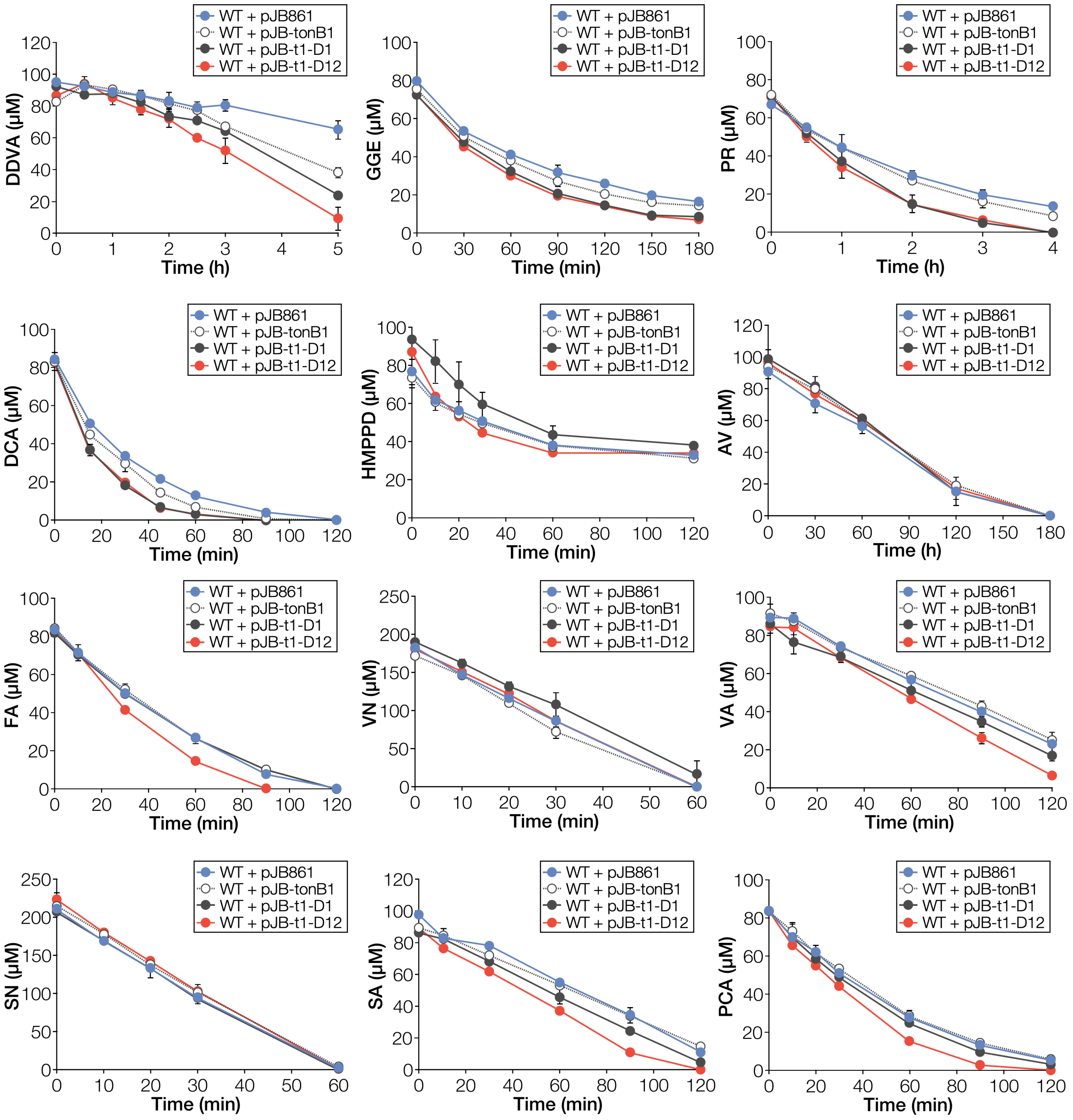


**Fig S9 Conversion of lignin-derived aromatic compounds by SYK-6 cells overexpressing the *tonB1* operon genes.** Cells of SYK-6(pJB861), SYK-6(pJB-tonB1), SYK-6(pJB-t1-D1), and SYK-6(pJB-t1-D12) were incubated with 100 µM DDVA, 100 µM GGE, 100 µM PR, 100 µM DCA, 100 µM HMPPD, 100 µM AV, 100 µM FA, 200 µM VN, 100 µM VA, 200 µM SN, 100 µM SA, and 100 µM PCA, respectively. Portions of the reaction mixtures were collected, and the amount of substrate was measured using HPLC. Each value is the average ± the standard deviation of three independent experiments.

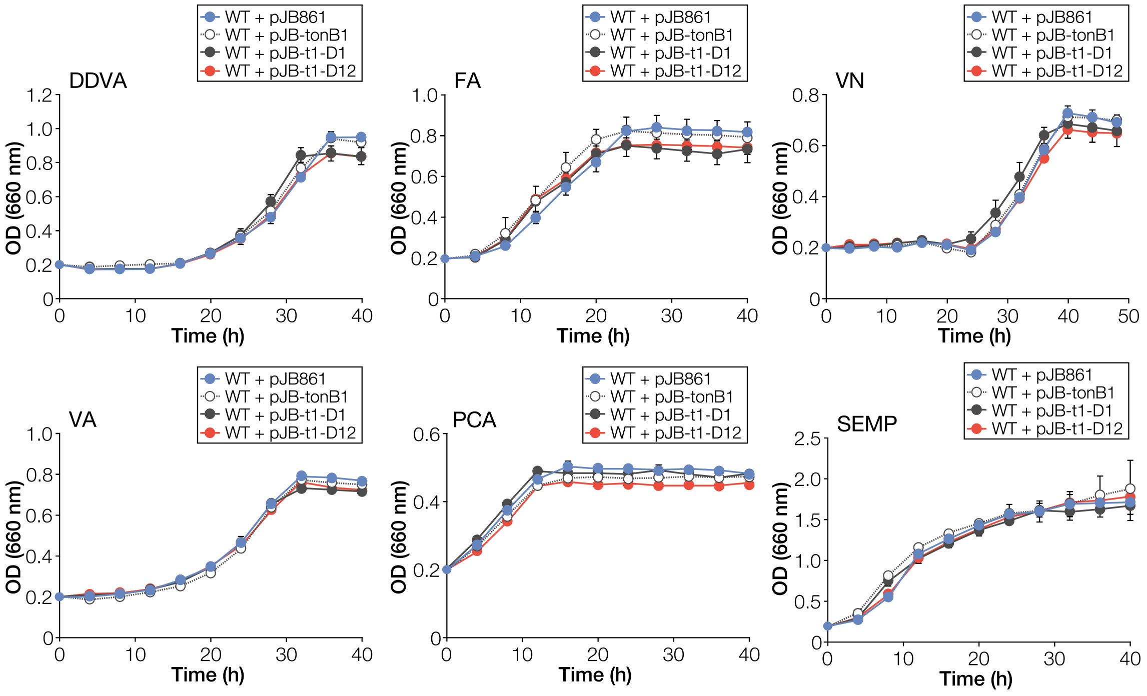


**Fig. S10 Growth of SYK-6 cells overexpressing the *tonB1* operon genes.** Cells of SYK-6(pJB861), SYK-6(pJB-tonB1), SYK-6(pJB-t1-D1), and SYK-6(pJB-t1-D12) were incubated in Wx medium containing 0.5 mM *m*-toluate and 5 mM DDVA, FA, VN, VA, or PCA and Wx medium containing 0.5 mM *m*-toluate and SEMP. Cell growth was monitored by measuring the $OD_{660}$. Each value is the average ± the standard deviation of three independent experiments.

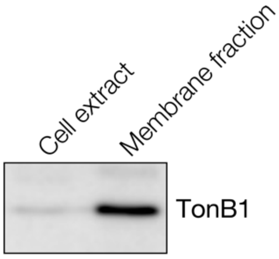


**Fig. S11 Cellular localization of TonB1.** Western blot analysis using anti-TonB1 antibodies was performed against the cell extract and total membrane fraction (10 µg protein each) obtained from SYK-6 cells grown in LB.

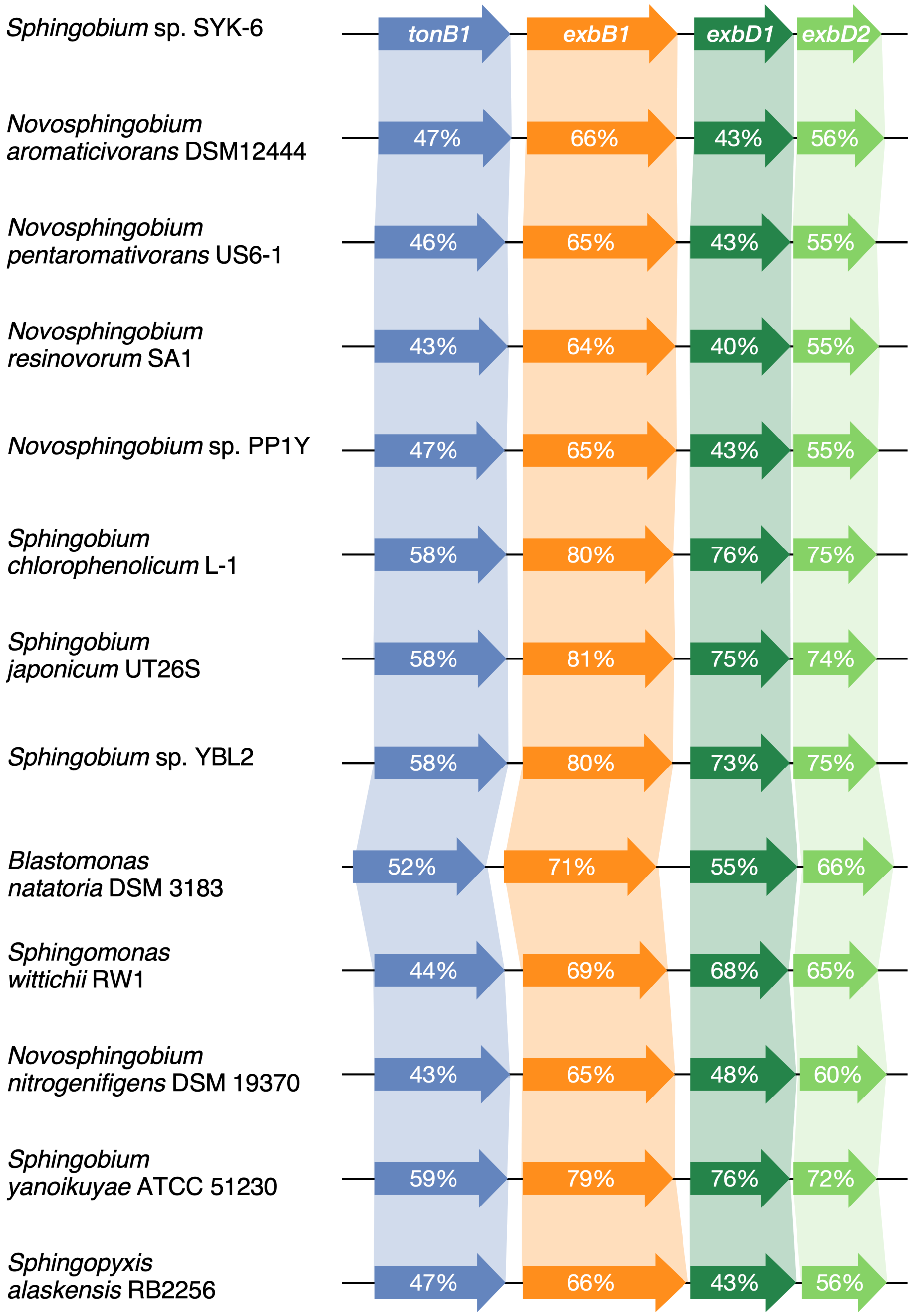


**Fig. S12 Organization of the Sphingomonadaceae genes showing similarity with the *tonB1* operon genes of *Sphingobium* sp. SYK-6.** Accession numbers of the genes are shown in Table S1.

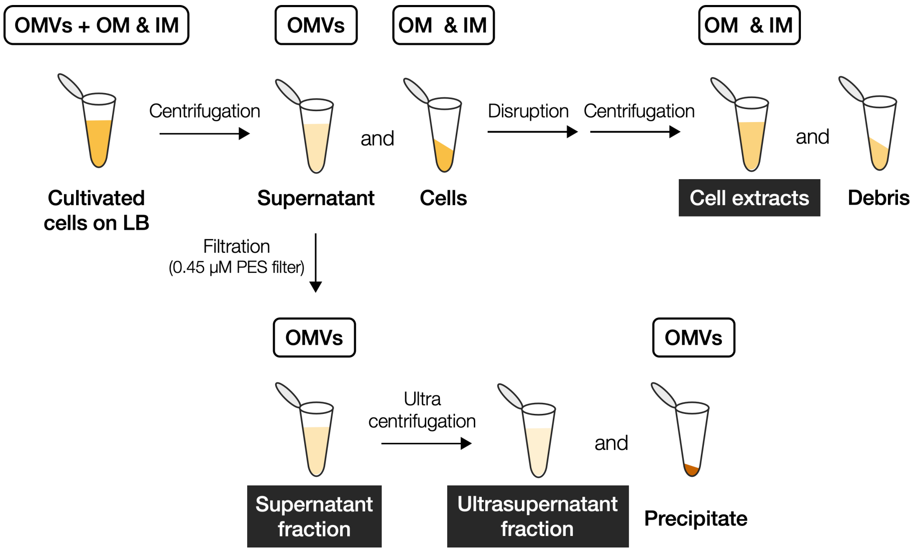


**Fig. S13 Scheme of sample preparation for western blot analysis in Fig. 6.** The supernatant fraction, ultrasupernatant fraction, and cell extracts were analyzed by western blotting. Details are given in the Methods. OMVs, outer membrane vesicles; OM, outer membrane; IM, inner membrane.

A

B

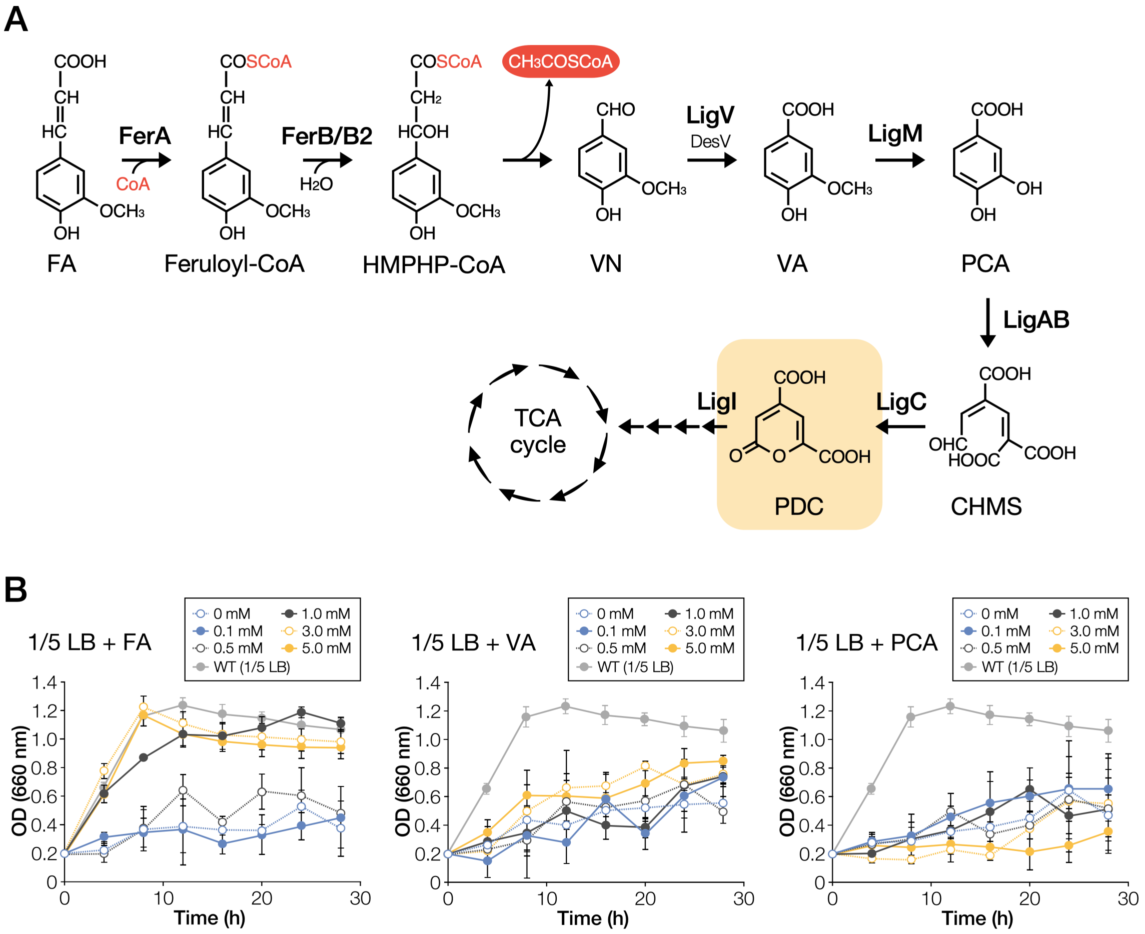


**Fig. S14 Addition of FA markedly improved the growth of Δ*exbD3*/*tolR ligI* in LB.** (A) Catabolic pathway of ferulate (FA) in SYK-6. Enzymes: FerA, feruloyl-CoA synthetase; FerB and FerB2, feruloyl-CoA hydratase/lyase; LigV, VN dehydrogenase; DesV, SN dehydrogenase; LigM, VA/3-*O*-methylgallate *O*-demethylase; LigA and B, small and large subunits of PCA 4,5-dioxygenase; LigC, CHMS dehydrogenase; LigI, PDC hydrolase. Compounds: HMPHP-CoA, 4-hydroxy-3-methoxyphenyl-β-hydroxypropionyl-CoA; CHMS, 4-carboxy-2-hydroxymuconate-6-semialdehyde; PDC, 2-pyrone-4,6-dicarboxylic acid. (B) Growth of Δ*exbD3*/*tolR ligI* cells in diluted LB with or without 0.1–5.0 mM FA, VA, or PCA. The growth of wild type cells in diluted LB shown in Fig. 2 is also indicated. Cell growth was monitored by measuring the $OD_{660}$. Each value is the average ± the standard deviation of three independent experiments.

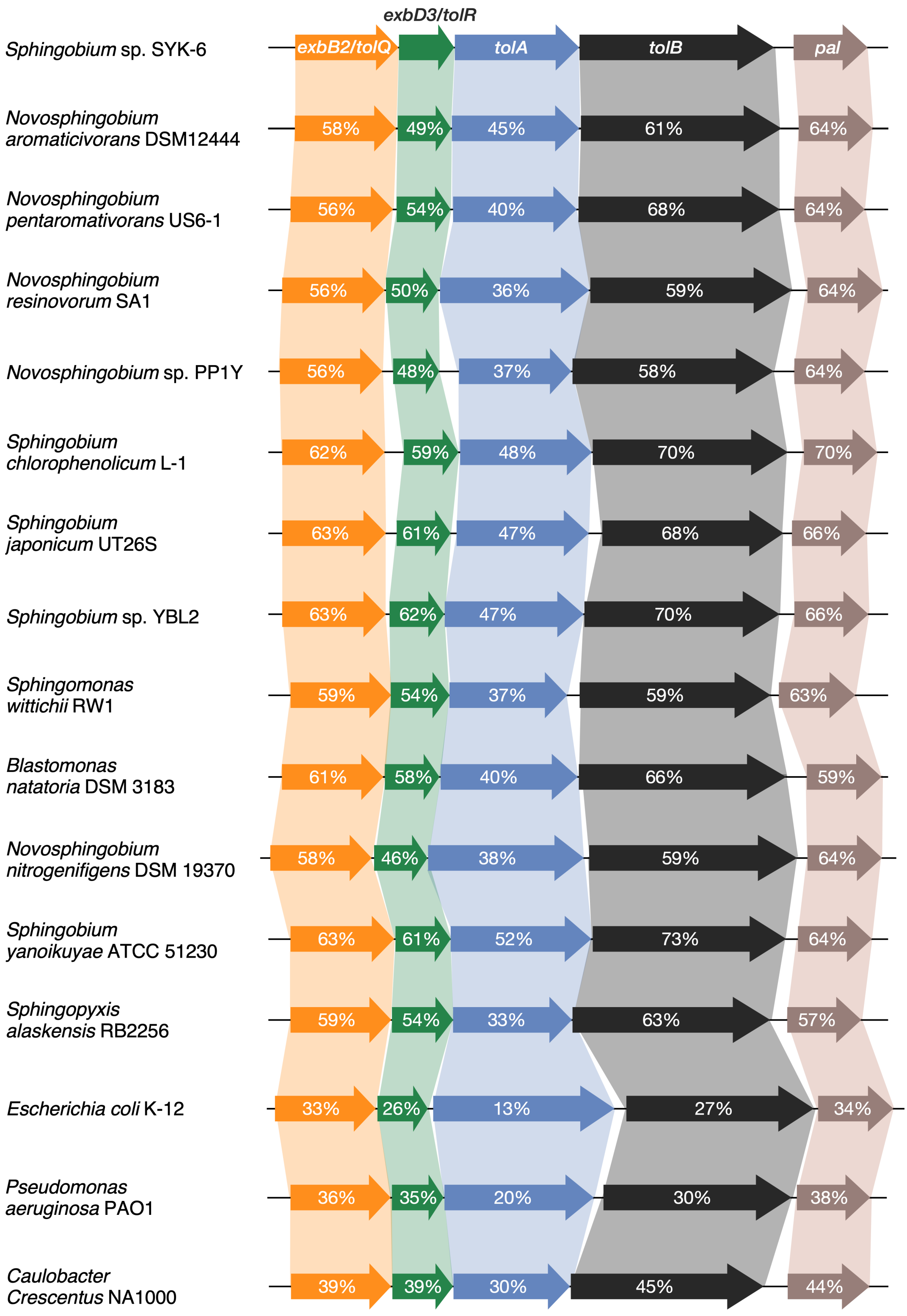


**Fig. S15 Organization of the Sphingomonadaceae genes showing similarity with the Tol-Pal system genes of *Sphingobium* sp. SYK-6.** Accession numbers of the genes are shown in Table S2.
